# Dynamic Routing on a Static Structure: Neuromodulatory Gating of Cortical Wave Propagation

**DOI:** 10.64898/2026.02.10.705210

**Authors:** Rishikesan Maran, James M. Shine, Ben D. Fulcher, Eli J. Müller

## Abstract

Stimulus-evoked cortical responses range from tightly localized activations to spatially extended traveling waves, yet the mechanisms through which these distinct regimes are flexibly recruited remain unclear. Based on neuroanatomical and neuroimaging evidence, we hypothesize that ascending neuromodulatory systems, which deliver modulatory (gain-changing) rather than driving inputs to cortex, dynamically shape spatiotemporal propagation under a fixed anatomical backbone. To test this hypothesis, we extend an empirically grounded corticothalamic neural field model to incorporate a diffusely projecting modulatory population, parameterized on the noradrenergic (NA) arousal system of the locus coeruleus (LC) as the best-characterized case. We then examine evoked responses under transient stimulus-locked versus sustained modulation—corresponding to phasic and tonic modulation - with geometry given by distance-dependent connectivity plus additional fast-conducting non-local projections (FNPs). Transient modulation produces spatiotemporally sustained traveling waves of cortical activity, whereas sustained modulation yields temporally prolonged but spatially localized responses; without modulation, waves are weak and dissipative. The model shows that the timing and spatial pattern of modulatory engagement gate wave-onset latency and impose anisotropy, restricting stimulus propagation to modulated territories and halting it across unmodulated gaps. In the presence of FNPs, which capture contributions of long-range white matter fiber tracts in the brain, modulation also enables the rapid non-local activation of distant cortical regions required for selective long-distance routing. Our results identify patterned gain control as a parsimonious mechanism for variability in evoked dynamics, and suggest that ascending modulatory systems such as the NA system could transiently privilege specific pathways over a largely static connectome.

**Author summary:** The brain doesn’t always react to a stimulus in the same way: sometimes the response is local, while at other times it sweeps across the entire cortical sheet as a wave. We propose that this variability can in part be controlled by ascending neuromodulatory systems which adjust the *gain* of cortical circuits rather than directly driving them. Using an empirically grounded corticothalamic model augmented with a modulatory population, we contrasted brief transient bursts versus sustained modulation. Transient modulation promoted robust traveling waves, while sustained modulation produced longer-lasting but spatially confined responses. By further specifying when and where it engages, modulation can gate where waves start, which directions they travel, and where they stop—even across unmodulated gaps. When long-range connections are present, modulation also enables rapid, selective activation of distant targets needed for long-distance routing. This neuromodulatory mechanism offers a simple, testable account of trial-to-trial variability in evoked dynamics, capturing a mechanism through which the brain can transiently privilege specific communication pathways through a static structural connectome.

## Introduction

The cerebral cortex exhibits broad variability in its responses to incoming sensory and externally applied stimuli(including magnetic, electrical, and optical), ranging from spatially localized activations to spatially diffuse traveling wave activations [1–6]. A key question is how this diversity of stimulus responses can arise from a structural connectivity that is relatively static on the timescale of the stimulus. A possible explanation to this phenomenon is a dynamic modulatory mechanism—complementing static connectivity—that can control the gain of excitability of constituent neural populations [7, 8]. Here, we explore whether ascending neuromodulatory systems, such as the noradrenergic system, can serve as a particular gain modulation mechanism that underpins dynamic variability in stimulus responses.

The propagation of stimulus-evoked activity through the axonal fibers of macroscale brain tissue has been observed in a range of experimental stimulation paradigms, including whisker deflections [6, 9], visual stimuli [10–12], transcranial magnetic stimulation [13, 14], and optogenetics [12, 15]. Importantly, cortical travelling wave dynamics have been observed in electrophysiological recordings [16, 17], as well as haemodynamic recordings [18–20], however, these occur over vastly different timescales and are measured via distinct biophysical processes.

In the cerebral cortex, fibers tend to be distributed more densely over shorter distances [21], causing propagation to be spatiotemporally organized as traveling waves of successively depolarizing populations [16, 22, 23]. There are also fibers that are distributed more sparsely over longer distances [24, 25] which can induce deviations from a wave dynamic by propagating punctate activity to remote populations, from which additional waves are initiated [9, 13, 15, 26]. Together, these corticocortical fibers of varying lengths constitute the structural (or anatomical) connectivity of the cerebral cortex, where its reorganization over an organism’s lifespan is negligible on the millisecond timescales of a single stimulus event [27, 28].

Although structural connectivity within the cerebral cortex is effectively static, experiments indicate that cortical neural response to repeated presentations of the same stimulus can exhibit substantial variability in spatiotemporal patterns between trials [1–6]. For example, whisker-deflection-evoked responses can exhibit sustained spreading processes (traveling waves and rapid long-range activations) in some trials, but can also exhibit weaker, spatially localized activations in others [6, 9]. Spreading variability has also been observed in the consciousness literature, where the sustained propagation of activity across the brain is termed ‘ignition’ and found concomitant with states of consciousness [3, 4, 29]. Furthermore, these spreading processes (when they occur) vary between trials in propagation direction [11, 23, 30–32] and latency of activation from stimulus onset [10]. Given that the structural connectivity of the cortex is relatively stable between trials, these studies therefore suggest that the brain’s evoked responses to stimuli are constrained not only by structural connectivity, but by an additional time-varying mechanism that mediates the extent to which the response spreads across the cortical surface.

Here, we propose spatially and temporally patterned gain modulation by the brain’s ascending neuromodulatory systems as a possible mechanism that can give rise to substantial diversity in evoked spatiotemporal responses of cortical activity to an identical input stimulus. This diversity includes localized versus spreading stimulus-evoked responses, and variability in wave propagation direction and latency. These systems share a common functional signature: at their target sites they can modulate the gain, i.e., the excitability of their target neurons, without necessarily eliciting an action potential [7, 33]. A population whose gain is modulated in this way may thereby control the variability between strong, spatially sustained responses under high gain, and weak, localized responses under low unmodulated gain.

For aspects of this study, we explore the noradrenergic (NA) system as our worked instantiation of this mechanism, as it is the most completely characterized of the ascending systems and therefore permits the tightest empirical parameterization. The NA system is a key brain system for promoting states of arousal and exploratory behavior [34], and comprises a range of nuclei in the locus coeruleus (LC), a structure located in the pons of brainstem, which project widely throughout the brain, including the cerebral cortex (Fig. 1A). While the LC collectively targets the cortex in a diffuse manner, the specific neurons of the LC that are activated, and the times of activation, vary significantly between trials [35, 36]. Based on these empirical observations, patterned modulation may also support spatiotemporal selectivity of activity propagation, resulting in cross-trial variability in spreading times and directions. We emphasize that the mechanism we characterize is general to modulatory gain control; the NA system supplies its parameterization and its empirical constraints, and we return in the Discussion to what would distinguish it from other ascending systems.

**Fig 1.**
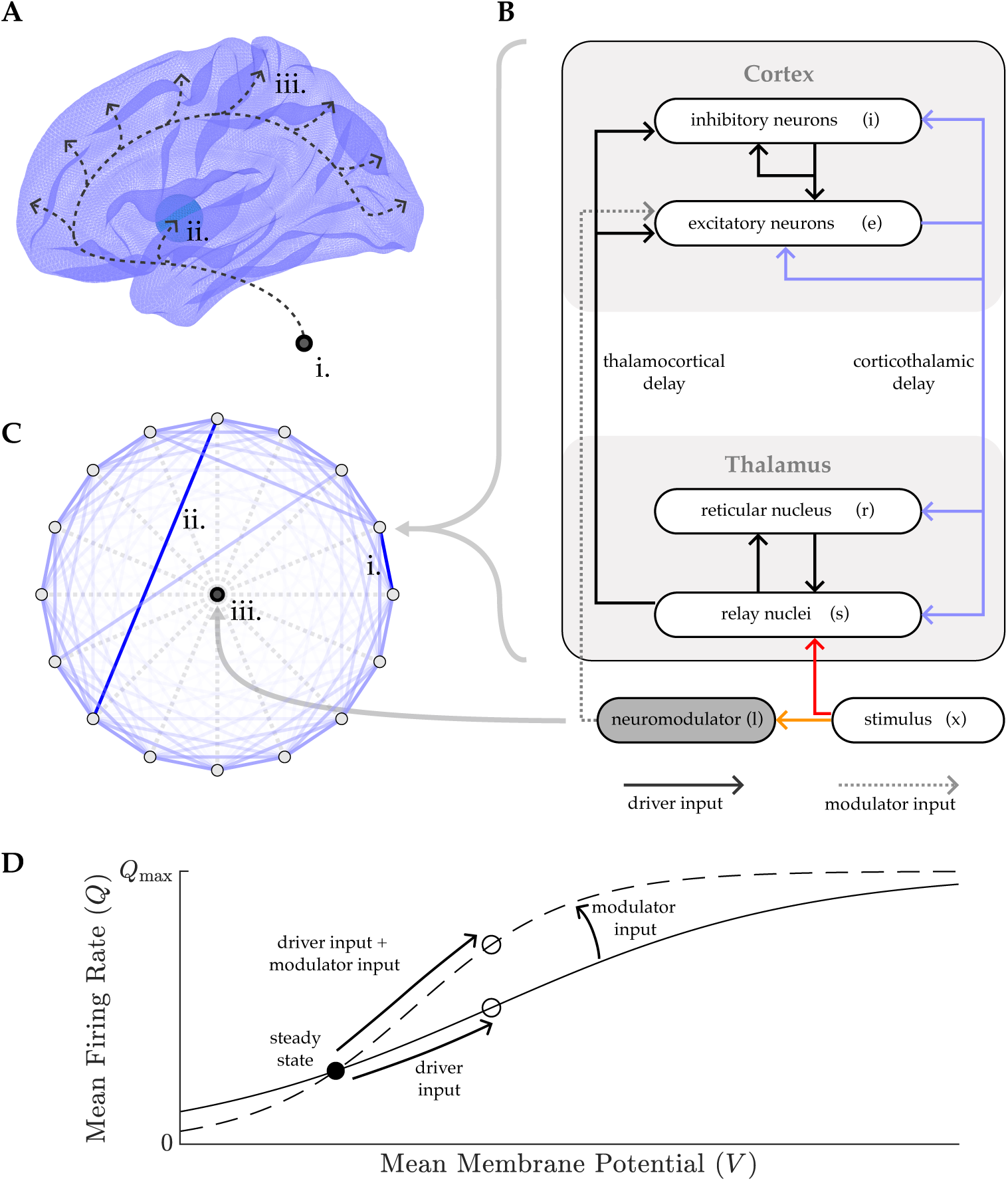
Schematic of corticothalamic model investigating the role of the ascending arousal system on macroscale corticothalamic dynamics. **A.** The **i.** neuromodulatory population projects to **ii.**) the thalamus, and **iii.** the cerebral cortex in a diffuse manner. **B.** The model tracks corticothalamic activity as a network of corticothalamic nodes. Each node tracks the mean activities of the excitatory (*e*) and inhibitory neurons (*i*) in distinct cortical regions, and the mean activities of the reticular (*r*) and relay (*s*) nuclei in their topographically aligned thalamic regions. **C.** The nodes are arranged into a circular ring, and are inter-coupled to each other, including **i.** stronger connections over shorter distances per the exponential distance rule (EDR), as well as **ii.** fast-conducting non-local projections (FNPs), which are long-range exceptions with faster conduction than other connections, and stronger connectivity than predicted by the EDR. The neuromodulator node (**iii.**) projects diffusely to all corticothalamic nodes. **D.** Each population *e, i, r, s* in each node has an activation function, which relates its mean membrane potential *V* to its mean firing rate *Q*. Driver inputs to a population increase its mean membrane potential, whereas modulator inputs increase its gain (the slope of the activation function about its steady state).

Understanding the contribution of the ascending arousal systems to the spatiotemporal properties of stimulus-evoked cortical dynamics is limited by the experimental difficulty of stimulating and recording neuromodulatory activity while simultaneously measuring its diffuse projection targets across the cerebral cortex.

Physiologically-based mathematical modeling offers a powerful solution to this barrier, by allowing the experimenter to simulate an approximation of neural activity both with and without the influence of neuromodulation. Several modeling approaches have sort to characterize this effect on cortical connectivity [37, 38], via cell-type specific interactions [29, 39], and dynamic routing in recurrent neural networks [40, 41] In addition, the resulting models can be used to elucidate how contributions, particularly from neuromodulatory structures such as the NA system, can shape, or are shaped by, other structural mechanisms within the brain, which may lead to specific hypotheses that can be tested in future empirical experiments.

In this study, we build on an empirically validated corticothalamic model, extending it to include an ascending diffuse neuromodulatory population - parameterized on the noradrenergic system - which is capable of modulating the gain of the existing system in the face of an incoming stimulus. Through a range of stimulus-response experiments, we show that variability in both the spatial targets and timing of neuromodulation yield corresponding variability in cortical responses to the same stimulus, where particular directions of activity propagation from the stimulus are prioritized over others. Taken together, our simulations demonstrate how patterned neuromodulatory gain control can support the brain’s diverse repertoire of spatiotemporal dynamics under a fixed connectivity structure. In addition, we demonstrate how timing via transient(phasic) gain modulation can both promote stimulus propagation along specific fibers whose endpoints lie within the modulated regions and, conversely, silence propagation along fibers that do not. This dynamic arousal mechanism could allow the brain to privilege some physical connections over others in a context-dependent manner, not only providing a mechanism for variability seen in stimulus-response experiments, but also a putative mechanism for how the brain may vary activity propagation streams vital for flexible, context-dependent cognition [42].

## Materials and methods

### Model description

To investigate the role of neuromodulatory systems on cortical activity, we extend a previous biophysical model of the corticothalamic system [43], to include input from an ascending neuromodulatory population (Fig. 1A-B). We provide a description of this novel model construction below, and detail the complementing model equations in S1 Appendix.

In the existing corticothalamic model, the cortex and thalamus are parcellated across space into a network of nodes, each of which has an architecture illustrated in Fig. 1B and C. Each node tracks the mean activities of excitatory (*e*) and inhibitory (*i*) populations in a distinct cortical region, and the mean activities of reticular (*r*) and relay (*s*) nuclei in the topographically aligned thalamic region. These populations can then project via axonal fibers to other populations in the same node, illustrated as solid arrows in Fig. 1B. The excitatory projections of each node, colored blue in Fig. 1B, can additionally project to other nodes. These excitatory projections have strengths which reflect the structural connectivity between the associated cortical regions, and time delays which reflect the finite velocity at which activity propagates axonally between the two regions. Such projections tend to be stronger over shorter ranges (Fig. 1C(i)), a phenomenon quantified by the exponential distance rule (EDR) [21, 44]. However, certain specifically positioned projections over longer distances exhibit strengths orders of magnitude larger than predicted by the EDR due to fiber bundling, and faster conduction velocities due to myelination (Fig. 1C(ii)) [24, 25, 45]. We call such projections fast-conducting non-local projections (FNPs). Note that the EDR also captures projections over long distances, but with much weaker connectivity strengths and slower conduction speeds.

Here we extend the existing corticothalamic model to include an additional single neuromodulatory population node (Fig. 1C(iii)), which projects to all existing corticothalamic nodes. These projections provide modulator inputs to the neural populations in other nodes (dotted lines in Fig. 1B), and are distinct from the existing driver inputs in the model (solid lines in Fig. 1B). As shown in Fig. 1D, the driving input to a population increases its mean membrane potential, which in turn increases its mean firing rate in accordance with its activation function. By contrast, the modulator input to a population increases its *gain*, which we quantify as the slope of its activation function about steady state [7, 46]. Modulator inputs hence have an indirect effect on a population’s mean firing rate, effectively amplifying the impact of any simultaneous driving input.

### Model Implementation

The parameters of the model and their values used in our investigations are given in Table 1. For clarity, we group the parameters based on the aspects of the model they govern: (i) the model topology: *N, R*; (ii) axonal propagation: *λ, γ, v, t*_0_; (iii) the dendritic response: *α, β, {ν_ab_}*; (iv) firing rates: *Q*_max_*, θ, σ*, and; (v) the modulation dynamics of population gain by the NM population: *η, {g_a_}, τ_el_*. Existing model parameters were previously fit to power spectra of human electrophysiological recordings in an ‘eyes-open’ state [47]. Below, we describe all newly incorporated parameters (highlighted bold in Table 1), and justify the choice of their assigned values.

**Table 1.**
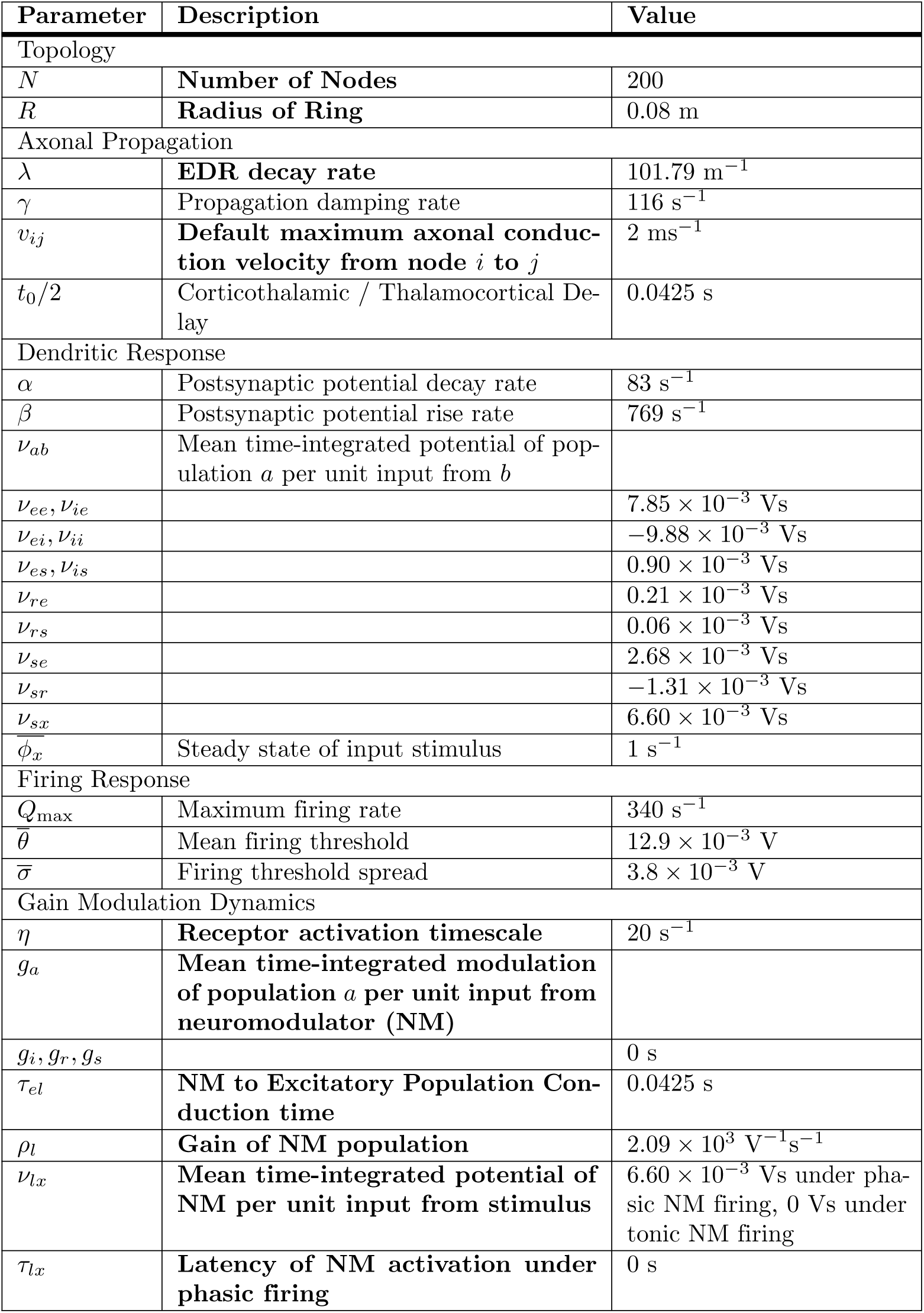
Nominal model parameter values for numerical experiments presented here. Newly incorporated parameters in the model are highlighted bold.

| Parameter | Description | Value |
|---|---|---|
| Topology |  |  |
| *N* | **Number of Nodes** | 200 |
| *R* | **Radius of Ring** | 0.08 m |
| Axonal Propagation |  |  |
| λ | **EDR decay rate** | 101.79 m<sup>−1</sup> |
| γ | Propagation damping rate | 116 s<sup>−1</sup> |
| *v*<sub>*ij*</sub> | **Default maximum axonal conduc-tion velocity from node *i* to *j*** | 2 ms<sup>−1</sup> |
| *t*<sub>0</sub>/2 | Corticothalamic / Thalamocortical De-lay | 0.0425 s |
| Dendritic Response |  |  |
| α | Postsynaptic potential decay rate | 83 s<sup>−1</sup> |
| β | Postsynaptic potential rise rate | 769 s<sup>−1</sup> |
| ν<sub>*ab*</sub> | Mean time-integrated potential of pop-ulation *a* per unit input from *b* |  |
| ν<sub>*ee*</sub>, ν<sub>*ie*</sub> |  | 7.85 × 10<sup>−3</sup> Vs |
| ν<sub>*ei*</sub>, ν<sub>*ii*</sub> |  | −9.88 × 10<sup>−3</sup> Vs |
| ν<sub>*es*</sub>, ν<sub>*is*</sub> |  | 0.90 × 10<sup>−3</sup> Vs |
| ν<sub>*re*</sub> |  | 0.21 × 10<sup>−3</sup> Vs |
| ν<sub>*rs*</sub> |  | 0.06 × 10<sup>−3</sup> Vs |
| ν<sub>*se*</sub> |  | 2.68 × 10<sup>−3</sup> Vs |
| ν<sub>*sr*</sub> |  | −1.31 × 10<sup>−3</sup> Vs |
| ν<sub>*sx*</sub> |  | 6.60 × 10<sup>−3</sup> Vs |
| φ̄<sub>*x*</sub> | Steady state of input stimulus | 1 s<sup>−1</sup> |
| Firing Response |  |  |
| *Q*<sub>max</sub> | Maximum firing rate | 340 s<sup>−1</sup> |
| θ̄ | Mean firing threshold | 12.9 × 10<sup>−3</sup> V |
| σ̄ | Firing threshold spread | 3.8 × 10<sup>−3</sup> V |
| Gain Modulation Dynamics |  |  |
| η | **Receptor activation timescale** | 20 s<sup>−1</sup> |
| *g*<sub>*a*</sub> | **Mean time-integrated modulation of population *a* per unit input from neuromodulator (NM)** |  |
| *g*<sub>*i*</sub>, *g*<sub>*r*</sub>, *g*<sub>*s*</sub> |  | 0 s |
| τ<sub>*el*</sub> | **NM to Excitatory Population Con-duction time** | 0.0425 s |
| ρ<sub>*l*</sub> | **Gain of NM population** | 2.09 × 10<sup>3</sup> V<sup>−1</sup>s<sup>−1</sup> |
| ν<sub>*lx*</sub> | **Mean time-integrated potential of NM per unit input from stimulus** | 6.60 × 10<sup>−3</sup> Vs under pha-sic NM firing, 0 Vs under tonic NM firing |
| τ<sub>*lx*</sub> | **Latency of NM activation under phasic firing** | 0 s |

To simplify model complexity in our investigations, the corticothalamic nodes are arranged with uniform spacing into a ring—a one-dimensional simplification of a single closed cortical hemisphere (Fig. 1C). This permits disassociation of explicit effects of spatiotemporal neuromodulation from contributions of more realistic, but more complex, brain connectivity geometries. In doing so, we can conveniently visualize the model’s traveling wave propagation over time, and isolate the roles of specifically arranged FNPs separately from the influence of geometry. In later sections, we demonstrate these principles can be readily translated to more realistic geometries using empirical estimates of human cortical connectivity.

We assigned a default geometric structural connectivity, where the connectivity strength between two nodes follows an EDR, i.e., decays exponentially with their distance. We fit this distance decay to empirical connectivity estimates described below. Internode distances were measured in two-dimensional space (instead of along the ring), consistent with the EDR in prior human brain modeling studies using distances in physical three-dimensional space (rather than along the cortical surface) [48].

We assigned the ring a total of *N* = 200 nodes, and set the ring radius *R* = 0.08 m, half the Euclidean length of the longest structural connections seen in human empirical connectivity estimates within the left hemisphere and which couple posterior to frontal cortical regions in these empirical data [19]. We fitted an EDR to these structural estimates within the left hemisphere using a nonlinear least-squares method to obtain an exponential decay rate of *λ* = 101.79 m*^−^*^1^ to 5 significant figures, consistent with previous literature [19]. The default model used throughout this work always includes the geometric constraint with connection strengths between all pairs of nodes (*j, k*) set to *c_jk_* = exp(*−λd_jk_*), where *d_jk_*is their Euclidean distance, and the maximum conduction velocity was set to *v_jk_* = 2 ms*^−^*^1^, following reported velocities of local traveling waves in the human cortex [16]. In addition, this falls within the reported estimates in macaques monkeys showing feedback/feedforward speeds between primary visual cortices *∼* 3.5 ms*^−^*^1^ and slower within area speeds *∼* 0.33 ms*^−^*^1^, and between middle temporal and primary visual areas *∼* 3 *−* 10 ms*^−^*^1^ [49]. However, in latter sections of this work, we explore the contributions of including FNPs in addition to the geometric connectivity, which increases both the connection strength, *c_jk_*, and conduction velocity, *v_jk_*, between the nodes connected via the FNP.

The gain modulation parameters of the model govern how the modulatory population shapes the gain of each cortical population over time. Although the present model and its interpretations are generally applicable to many ascending neuromodulatory systems, for clarity we constrain our present model by parameterizing some of these neuromodulatory properties from empirical works on the NA system as a specific case study, as follows. At the microscale, LC neurons send noradrenaline to the adrenoceptors of target neurons [50], which in turn alters their firing thresholds [51]. Aggregating this microscale process over a population, we assumed that the change in each target population’s gain is proportional to the change in concentration of activated adrenoceptors, implying that the rate at which the gain changes is equal to the rate of adrenoceptor activation [7]. We set this rate to *η* = 20 s*^−^*^1^, following reports that the timescale of adrenoceptor activation is approximately 0.05 s [52]. Analogous metabotropic receptor kinetics govern the other ascending modulatory systems, though with system-specific timescales. *g_a_* is the strength at which population *a* is modulated. Although the LC as an aggregate has widespread projections across the brain [53], our aim was to investigate the effect of modulation on the spatial propagation of activity over the cerebral cortex, which is in turn facilitated by the excitatory propagators of each node. Thus, we allowed only *g* = *g_e_* to change, and fixed *g_i_* = *g_r_* = *g_s_* = 0, i.e., only cortical excitatory gain is modulated. Finally, since ascending neuromodulatory systems are subcortical, we assumed that the conduction time from the NM to excitatory populations is equal to the thalamocortical delay (*τ_el_* = *t*_0_*/*2), and that the gain of the NM is equal to the steady-state gain of the relay population (*ρ_l_* = *ρ_s_*).

To simulate the model’s stimulus-evoked response, we set the stimulus propagator, denoted by the red arrow in Fig. 1B, as an impulse (delta function). This input was fed to the relay population of one node of the ring to mimic a sensory-like stimulus that reaches a single cortical region via the thalamus, such as the input of a visual stimulus to primary visual cortex [29], or a whisker stimulation input to the corresponding barrel cortex [6].

Again, although our model is generalizable to many ascending arousal systems, we focus here on the noradrenergic system to constrain some of its parameters, as these has been characterized well empirically. The LC receives afferents from many brain regions, including the medial prefrontal cortex and hypothalamus [53], empirical observations have shown phasic activations of the LC that are time-locked to task specific stimuli during behavior [35, 54]. Here, we approximate this relationship between phasic firing behavior of neuromodulatory systems and stimuli, by coupling the stimulus to the NM population (*τ_lx_* = 0), denoted by the orange arrow in Fig. 1B. We then go on to explore the effect of imposing a lagged temporal relationship as part of our investigation. This coupling has the same synaptic strength as the relay population of the stimulus node (*ν_lx_* = *ν_sx_*), consistent with the model’s random connectivity assumption [47]. Finally, to model tonic modulation, the NM population is decoupled from the stimulus (*ν_lx_* = 0), and set to a constant firing rate.

All model equations were solved numerically using a finite difference scheme with timestep Δ*t* = 2*^−^*^14^ s. Second-order ordinary differential equations were solved using centered finite difference methods, while first-order equations were solved using the ‘Backward Euler’ method [55]. Details about these numerical schemes are in S2 Appendix.

## Results

Using our novel neuromodulated corticothalamic model, we investigate how modulating cortical gain through ascending arousal systems—tuned by strength, cortical targets, and timing—can affect the spatiotemporal properties of the observed stimulus-evoked responses. In addition, we explore how this dynamic gain modulation impacts the contributions of long-range connectivity (FNPs) to the spatiotemporal cortical activity dynamics.

### Effect of phasic and tonic modulation on stimulus-evoked dynamics

Ascending neuromodulatory systems commonly exhibit two distinct firing modes: (i) phasic, in which the modulatory population transiently increases its firing rate in response to a task-relevant stimulus; and (ii) tonic, in which it maintains an increased constant firing rate uncoupled from the stimulus. Both modes are well characterized for the LC [56, 57], whose parameterization we adopt here.

We investigate how the model’s stimulus-evoked cortical response, denoted by variable *ϕ^j^*(*t*) (S1 Appendix), is shaped uniquely by each modulation mechanism. For these experiments, we fixed a default geometric structural connectivity (a fully connected network with edge weights set by the EDR) and, in the presence of phasic modulation, set the NM population to fire time-locked to the stimulus - although we explore this temporal relationship in the following section.

We first examined the stimulus-evoked response when there was no modulation mechanism, indicated by the absence of the NM population (Fig. 2A(i)). Figure 2A(ii) illustrates the evoked response (*ϕ^j^*(*t*) for each node *j* = 0*, . . . ,* 199 over time *t*) from the time of stimulus onset, when the impulse stimulus was applied to node 100 (illustrated by the red dot on the figure’s vertical axis). We also plot the modulation factor over time, which is the gain of the excitatory population divided by its pre-stimulus (unmodulated) value. The evoked response takes the form of a highly dissipative traveling wave emanating from the stimulated node. This traveling wave rebounds approximately every *t*_0_ = 85 ms, which arises from activity propagating to the thalamus and back to the cortex via the corticothalamic loop resonance [43]. The peak activation (maximum activity) of the traveling wave decays rapidly across the cortical sheet with distance from the stimulated node due to dissipation (Fig. 2A(iii)).

**Fig 2.**
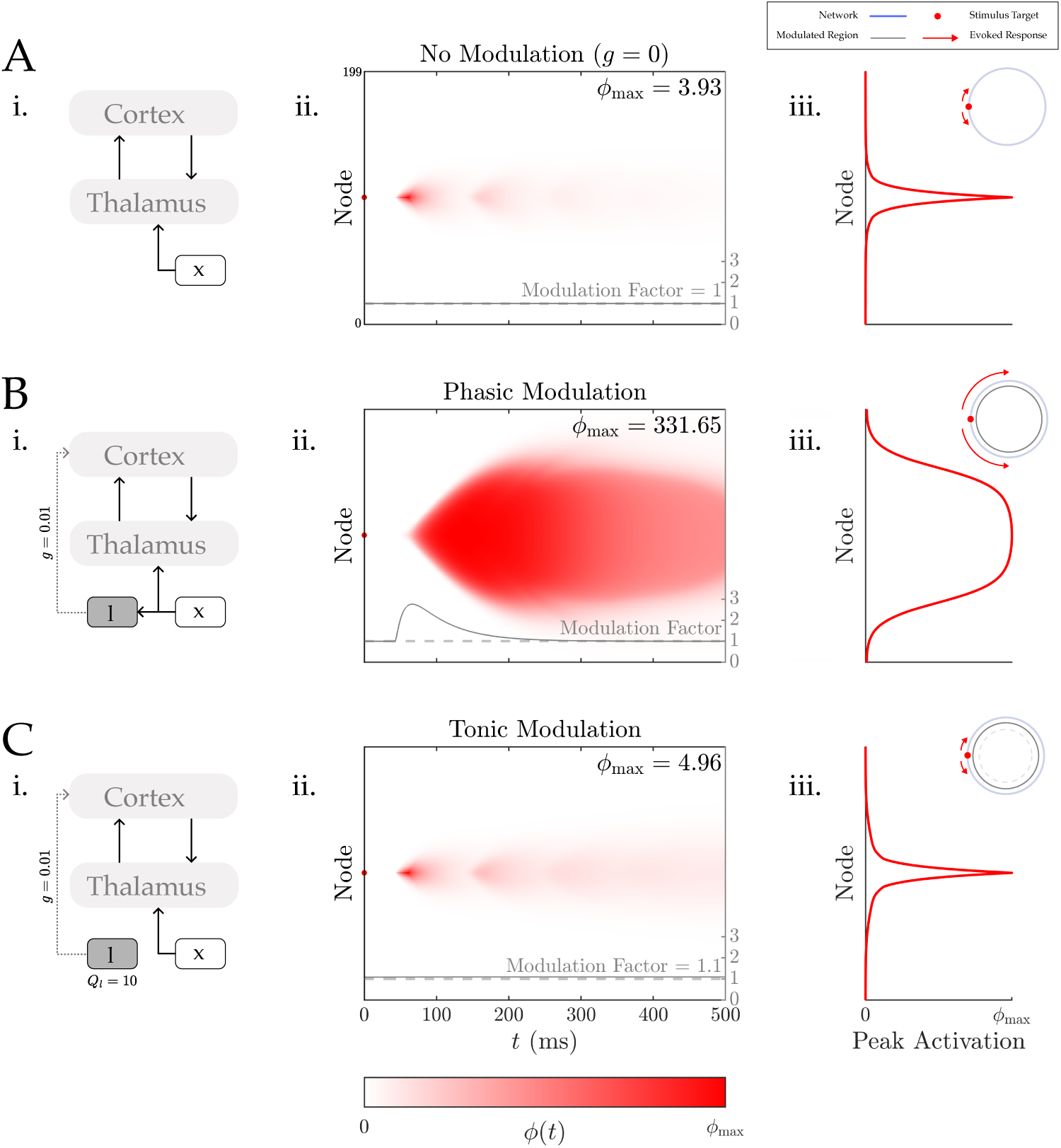
Phasic modulation shapes stimulus-evoked responses by enabling spatiotemporally sustained cortical traveling waves. We examined the effect of phasic and tonic modulation on the model’s stimulus-evoked dynamics, by comparing the response to the same stimulus under three different NM system configurations: **A.** NM was absent, so no modulation took place; **B.** The NM population fires phasically in response to the stimulus, and modulates the cortex with strength *g* = 0.01 s; **C.** The NM population fires tonically at a constant firing rate *Q_l_* = 10 s*^−^*^1^, uncoupled from the stimulus, and modulates the cortex with strength *g* = 0.01 s. The nodes were numbered from 0 to 199 clockwise on the ring, and the stimulus was applied to node 100. For all three configurations: **i.** the wiring configuration between populations in each node; **ii.** heatmaps of the stimulus-evoked response (*ϕ^j^* (*t*) (s*^−^*^1^) for each node *j* = 0*, . . . ,* 199 over time *t*) and the modulation factor, both from the time of stimulus onset; and **iii.** the peak activation of each node *j* (the maximum attained value of *ϕ^j^* (*t*) over *t*). In **ii.**, the top-left inset depicts all nodes arranged in a ring (blue), the stimulus node (red) and an indication of modulation applied to the nodes (gray). In **ii.** and **iii.**, *ϕ*max is the maximum attained value of *ϕ^j^* (*t*) across *j*, *t*.

Next, the effect of a phasic modulation mechanism was investigated by coupling the stimulus to the NM (population *l*), which in turn modulated the excitatory cortical population with strength *g*= 0.01 s (Fig. 2B(i)).

Previous work has mapped the stability properties of this corticothalamic model as a function of cortical gain, and how these aggregate to define a linear stability manifold [43, 47]. As the modulation here occurs exclusively for the excitatory cortical population, this increases the linear gains for each output of the cortical excitatory population and drive the system towards an instability, i.e., amplifying the evoked response through the corticothalamic loop [58] for each node.

The network evoked response under this configuration (Fig. 2B(ii)) is more sustained over time and propagates over a larger proportion of the ring. Figure 2B(iii) shows that although peak activation is still maximum at the stimulus node, this decays more slowly with distance from this node. We note, the high-firing rates are due to the multi-stability properties of the model for this parameter configuration [47] and the presence of a high firing rate attractor. The present study focuses on the first-phase evoked responses and their spatial organization by normalizing the stimulus drive to the thalamus and thus firing rate changes following stimulation should be taken as qualitative in the absence of scaling to physiological bounds.

Finally, we investigated how tonic modulation impacts evoked responses. As illustrated in Fig. 2C(i), the mean firing rate of the NM population was set to *Q_l_* = 10 s*^−^*^1^, and was closest to the largest possible firing rate at which the steady state of the model remained dynamically stable (results for higher firing rates can be found in S1 Fig). We found that tonic modulation at levels maintaining linear stability caused prolonged evoked responses temporally, but without extending it spatially (Fig. 2C(ii)). Previous work has shown that sustained diffuse projections - providing first order effects of neuromodulatory gain regulation - can act as a control parameter reducing the distance to a bifurcation and thereby temporally extending stimulus evoked responses [59]. We find the same effect here, and additionally show that at neuromodulatory levels consistent with system stability, tonic modulation supports spatially localized evoked responses (Fig. 2C(iii)). We note, this is necessarily a feature of the level of tonic modulation possible in the stable regime and not a feature of tonic modulation more generally.

In summary, the model’s traveling wave response to an impulse stimulus is relatively weak and localized in the absence of neuromodulation, strong and spatiotemporally sustained under phasic modulation, and temporally sustained but spatially localized under tonic modulation supporting stable dynamics. These results support phasic modulation as a candidate neurobiological mechanism for facilitating spatially sustained traveling wave responses on short timescales, while maintaining system stability over longer timescales.

### Effect of modulation timing and cortical targets on evoked dynamics

In the previous section we explored the effects of phasic neuromodulation with a fixed timing relationship to the stimulus onset and, importantly, a uniform spatial modulation of gain, i.e., all cortical nodes were targeted. However, neuromodulatory systems can show significant heterogeneities - in particular individual LC neurons show variable phasic activation relative to stimulus onset in empirical recordings across repeated trials [35, 36, 60, 61], but spatially sustained traveling wave responses also vary in their timing relative to stimulus onset [10]. In addition, neurons of the LC show substantial heterogeneity in their projection profiles to the cerebral cortex, targeting unique cortical regions, as well as a predominance of asynchronous activity between the neurons [62]. Together, these key features align with empirically observed variability of the directions of propagation of traveling wave responses [11, 30, 31]. We next investigated whether these empirical spatiotemporal variations in stimulus response can be captured by adjusting the relative timing of neuromodulatory firing and spatial distribution of its active cortical projections.

First, we examined how the timing of neuromodulation relative to stimulus onset affected the temporal properties of the stimulus-evoked response. We introduced a delay between the time of NM phasic firing and stimulus onset (parameter *τ_lx_*). As shown in Figs 3A(i) and A(ii), delays of *τ_lx_*= 25 ms and 50 ms resulted in corresponding delays in the time at which the wave begins to travel from the stimulus node, despite a fixed stimulus onset time. However, the peak activation of each node also tends to reduce with increasing delays, and with sufficiently large delays of *τ_lx_ >* 400 ms the modulator input eventually misses the stimulus with peak response more than 10-fold smaller S2 Fig. This is consistent with empirically observed timescales of LC influence on cortical activity. This result suggests that empirically observed variability in the timing of phasic neuromodulation can contribute to variable timing in traveling waves of evoked responses, despite receiving an identical stimulus.

**Fig 3.**
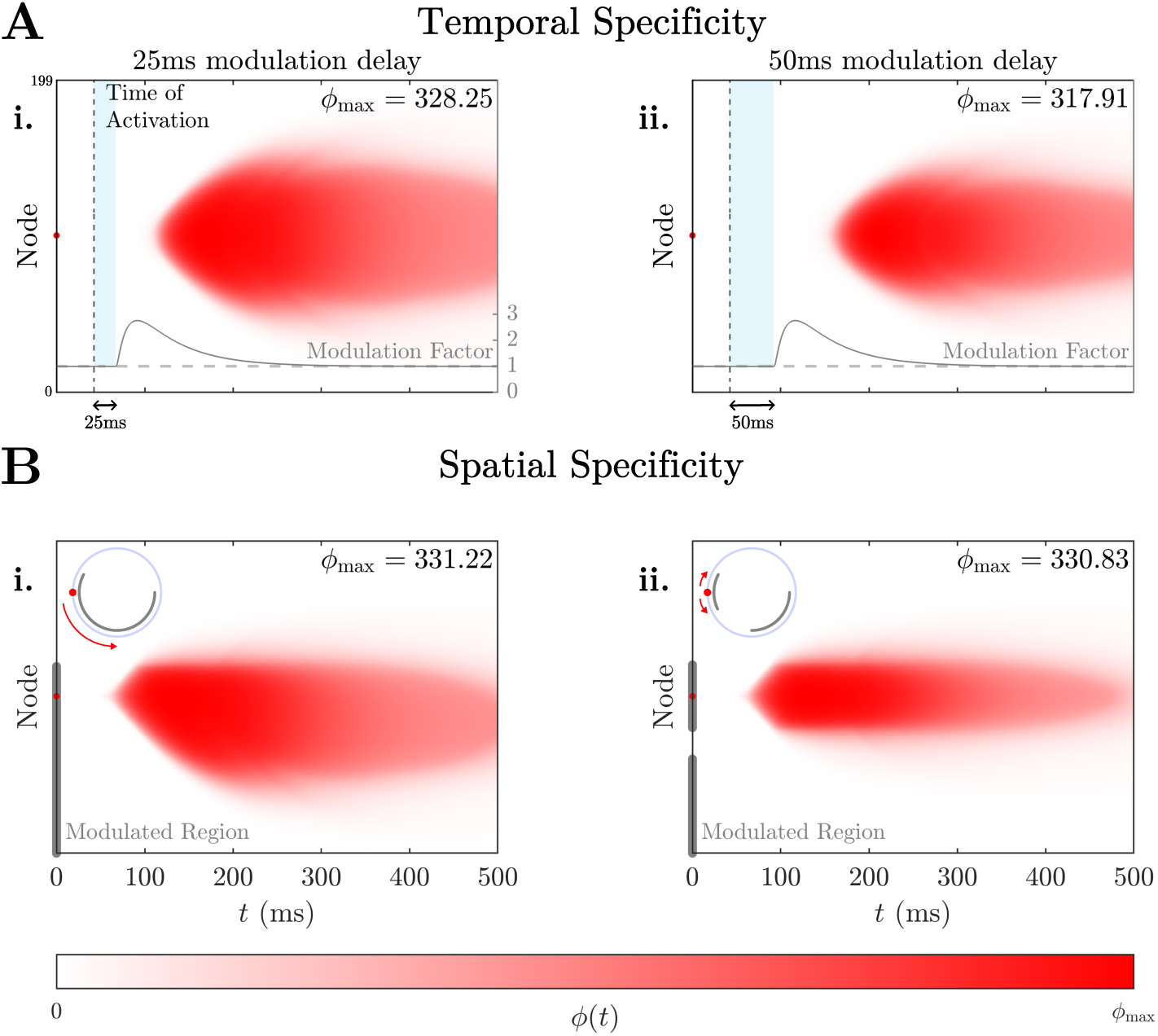
Temporal and spatial specificity of neuromodulation of the cortex shapes spatiotemporal evoked cortical traveling waves. **A.** We examine how the timing of modulation relative to stimulus onset affected the timing of evoked traveling waves. Heatmaps of the stimulus-evoked responses and the modulation factor over time are illustrated when the time of NM firing (relative to stimulus onset) was delayed by: **i.** 25 ms; and **ii.** 50 ms. **B.** Impact of restricting the cortical targets of the NM to a subset of cortical nodes would affect the propagation of a cortical traveling wave. The modulated region of nodes is highlighted gray on the vertical axis. For all four simulations, the modulation strength of the cortex was *g* = 0.01 s. The stimulus location is shown with a red dot (node 100) on the vertical axis of all plots, as is the maximum response of any node, *ϕ*max is annotated.

Next, we explored how varying the set of specific nodes that the neuromodulation projects to can give rise to spatial bias in the direction of propagation of the evoked response. In Fig. 3B(i), we restricted the set of nodes that were targeted by gain modulation to the set *{*0, 1*, . . . ,* 120*}*. We find that, compared to the simulated response when all nodes were modulated (Fig. 2B(ii)), the evoked traveling wave is restricted to the modulated region, i.e., it stops propagating at the boundary of this region. Hence, despite isotropic corticocortical structural connectivity, the spatially nonuniform distribution of neuromodulatory projections to the cortex introduces anisotropy to the propagation of traveling waves. Specifically, activity propagates preferentially through modulated cortical areas, capturing a key observation found in experimental animal studies [30, 31]. Furthermore, as depicted in Fig. 3B(ii), when the modulated region was *{*0, 1*, . . . ,* 60*} ∪ {*80*, . . . ,* 120*}*, so that a spatial gap is introduced, the evoked traveling wave fails to propagate across this gap, and instead dissipates at the boundaries of the modulated patches. Hence, the spatial contiguity of neuromodulation is an additional determinant of the extent to which wave propagation is sustained in space. Conversely, disjointedness between modulated regions, which may result from the regional specificity of neuromodulatory projection distributions to the cortex [62, 63], can give rise to compression of evoked traveling waves, which have been observed empirically at boundaries between cortical regions [11].

### Effect of gain modulation on long-range cortical propagation

In addition to traveling waves, spreading processes during stimulus-evoked responses can also occur along fast-conducting non-local projections (FNPs) between remote cortical populations [6, 9]. These rapid propagations can be interpreted as a perturbation of the traveling wave dynamics, by introducing non-local pairwise interactions that cannot be accounted for by the local cortical geometry [48, 64]. Here, we test whether gain modulation can explain this non-local perturbation during stimulus-evoked responses, by introducing a single FNP to the existing geometric structural connectivity. We set the FNP to bidirectionally connect nodes 50 and 150, so that its length (*d*_50,150_ = 2*R* = 0.16 m) is comparable to empirical estimates of the longest intra-hemispheric white-matter fibers in human (highlighted green in Fig. 6A) which couples posterior and frontal cortical regions (visualized in Fig. 6B). Starting from the geometric connectivity strength after normalization, the strength of this bidirectional FNP was increased by a factor of 10^5^. This aligns with the average normalized strength of empirical estimates of the posterior-frontal fibers (8.9 *×* 10*^−^*^4^), which was approximately a factor of 10^5^ times the normalized strength of the projection between the two nodes under geometric connectivity (exp(*−λ · d*_50,150_)*/*^199^ exp(*−λ · d*_50*,j*_) *≈* 10*^−^*^8^). We set the minimum conduction time of the FNP to the idealized case of 0 s. This approximation allows us to investigate the qualitative perturbatory effect of a non-local interaction (on a timescale much faster than that of geometric propagation) on the model dynamics. The zero-delay case is a deliberate limiting case that maximally separates FNP-mediated from geometric propagation in time and therefore yields an upper bound on the perturbation a single FNP can induce; this will necessarily place a lower bound on the modulation required for the FNP to become consequential. We revisit this assumption later in the manuscript.

An impulse stimulus was applied to node 50, an endpoint node of the FNP, to maximize the perturbation that the FNP induces on the consequent dynamics (labeled red in Figs 4A,B). We then compared the response between two structural connectivity configurations: geometric connectivity without the FNP (top-left inset of Fig. 4A(i)); and perturbed connectivity containing the additional FNP (top-left inset of Fig. 4B(i)).

**Fig 4.**
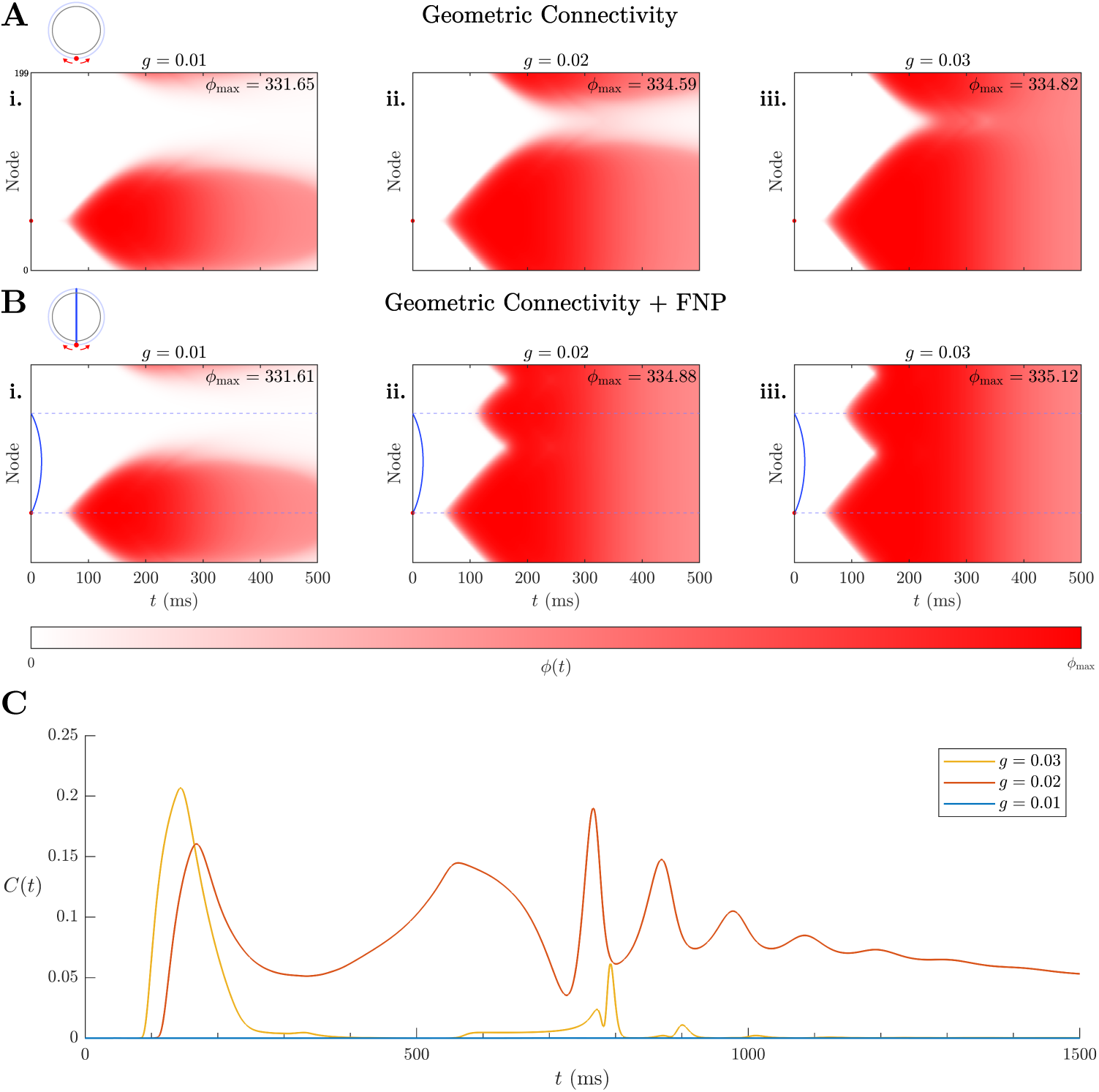
Neuromodulation enables rapid non-local activity propagation facilitated by fast-conducting non-local projections (FNPs). We examined the influence of a single FNP on stimulus-evoked dynamics for different modulation strengths, by comparing the evoked dynamics under two structural connectivity configurations. **A.**Under default geometric connectivity (top-left inset of **i.**), the evoked response to an impulse stimulus at node 50 is simulated and shown over time for three modulation strengths: **i.** *g* = 0.01, **ii.** *g* = 0.02, and **iii.** *g* = 0.03 s. **B.** Under an alternative connectivity that additional contains a bidirectional FNP between nodes 50 and 150 (top-left inset of **i.**), the evoked response to an impulse stimulus at node 50 is simulated and shown over time for the same three modulation strengths. The FNP has connectivity strength increased by a factor 10^5^, and minimum conduction time of 0 s. **C.**For each *g*, the cosine dissimilarity (or cosine distance) between the spatial activity patterns of the geometric and perturbed dynamics over time, denoted *C*(*t*) (Eq. (**??**) in S3 Appendix), quantifies the perturbation to the simulated dynamics caused by the addition of the FNP, relative to the geometric dynamics.

Figures 4A(i) and B(i) indicate that when all *N* nodes are modulated with strength *g* = 0.01 s, the evoked responses to the stimulus are similar under both connectivities, i.e., the added FNP minimally perturbs the dynamics in this uniform low-modulation setting. When the modulation strength is increased to *g* = 0.02 s or *g* = 0.03 s (Figs 4A(ii-iii) and B(ii-iii)), the FNP perturbs the dynamics more noticeably, initiating an additional traveling wave from node 150, the other endpoint of the FNP, at time *t* ≳ 100 ms. To quantify the FNP’s perturbation on the evoked dynamics for each value of *g*, we quantified the cosine dissimilarity (or cosine distance) between the spatial activity patterns of the geometric and FNP perturbed responses over time *t*, denoted as *C*(*t*) (see Eq. (**??**) in S3 Appendix for formulation) [64]. A plot of *C*(*t*) in Fig. 4C shows that the FNP’s perturbation remains minimal for *g* = 0.01 s, but deviates from zero for *g* = 0.02, 0.03 s at time *t ≈* 100 ms, which closely corresponds to the time at which node 150 activates. These results demonstrate that neuromodulation enables not only sustained traveling wave activity propagation via geometric connectivity, but also rapid non-local propagation via spatially heterogeneously distributed FNPs, and whose effects are dominant on short timescales - as shown in previous work [64].

Figure 4C shows that the FNP-induced perturbation, *C*(*t*), exhibits a sharp transition from *C*(*t*) *≈* 0 at *g* = 0.01 s (i.e., the FNP has a negligible effect on the dynamics), to non-trivial *C*(*t*) *>* 0 dynamics at *g* = 0.02 s (i.e., the FNP substantially perturbs the model dynamics), suggesting a threshold-like dependence on modulation strength. To systematically characterize this property, we examined how the temporal dynamics of *C*(*t*) varies across a more finely resolved spectrum of *g* values. Figure 5A shows that *C*(*t*) sharply deviates from near-zero behavior at a threshold *g ≈* 0.015 s. Above this threshold, modulation amplifies activity at the FNP’s distal endpoint to detectable levels before the traveling wave arrives; below this threshold, the FNP continues to propagate activity but at levels too small to distinguish from traveling wave dynamics under geometric connectivity alone. This sharp, threshold-like dependence of the functional impact of an FNP on neuromodulation strength suggests that the detectability of FNP-mediated non-local propagations depends critically on whether modulation supplied to the cortex exceeds some minimum level.

**Fig 5.**
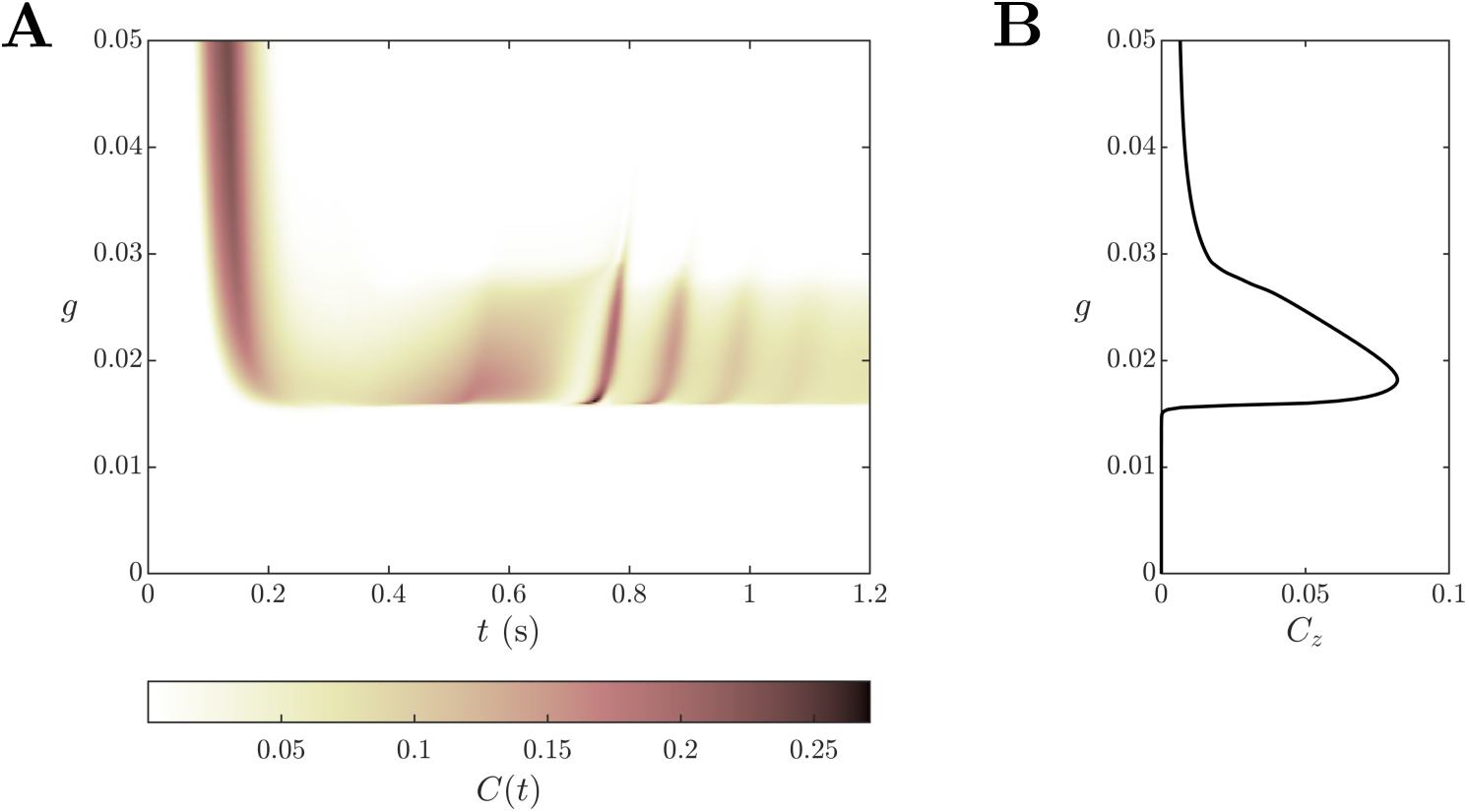
Moderate neuromodulation maximizes effects of long-range connections. **A.** The perturbation of the FNP between nodes 50 and 150 on the evoked response to a stimulus at node 50, measured as *C*(*t*), is plotted over time for a finely resolved set of modulation strengths *g*. **B.** For each *g*, the cosine dissimilarity between the zero-frequency components of the geometric and perturbed response patterns, denoted *Cz* (Eq. (**??**) in S3 Appendix), quantifies the FNP’s perturbation on the evoked dynamics (relative to geometric connectivity) over long timescales, *t >* 100 ms.

**Fig 6.**
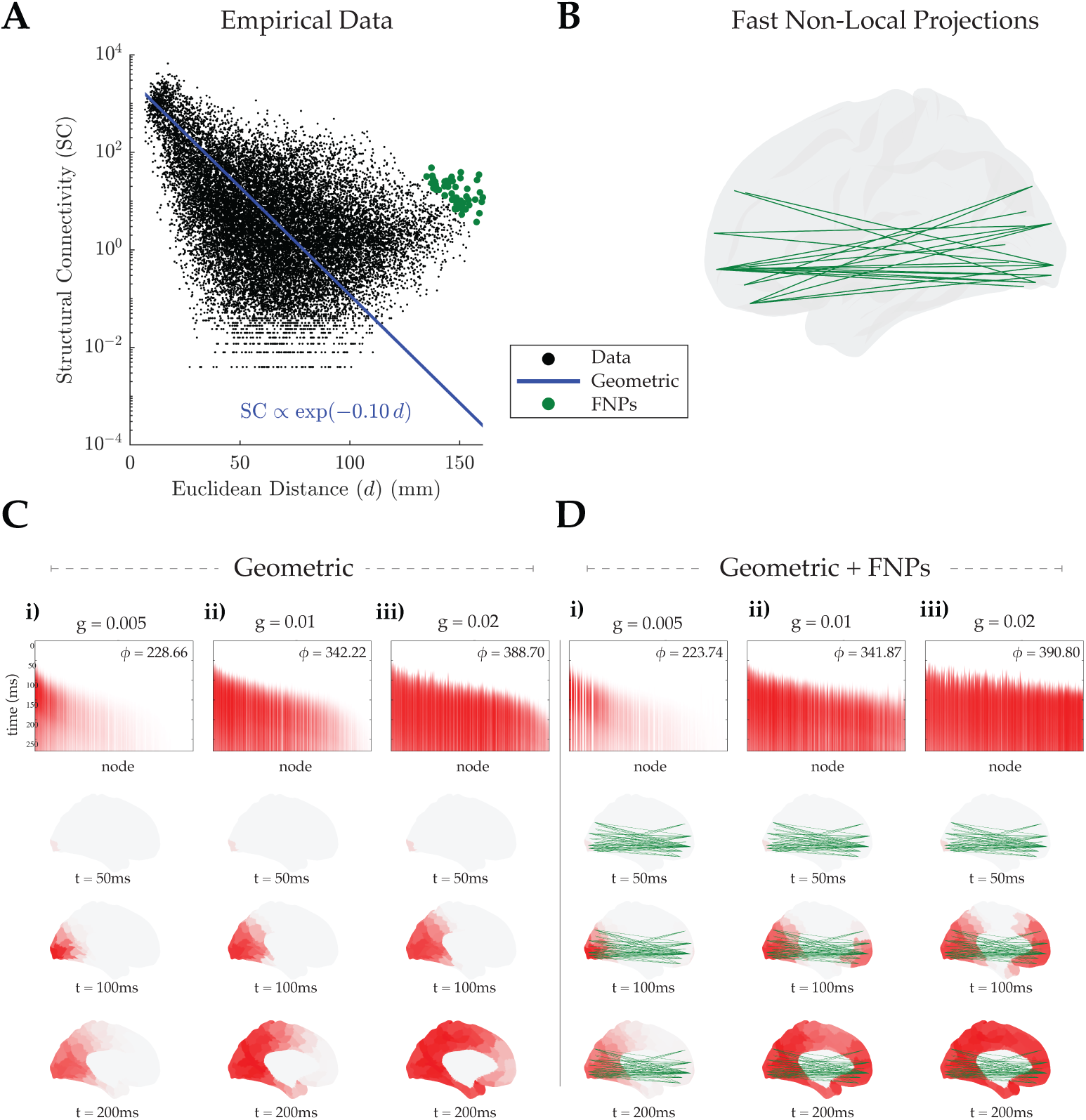
Empirical human structural geometry. **A.** We attained the high-resolution connectivity map from diffusion MRI tractography [19] giving pairwise estimates of the structural connectivity between the 29, 696 vertices of FreeSurfer’s ’fsaverage’ population-averaged template that are outside the medial wall. We then created a parcellated version (Schaefer-400 [66]) of this connectome, in which the structural connectivity between each pair of cortical regions is the weighted sum of all vertex-vertex connections that lie between the two pairs. Then, the Euclidean distance between each pair of regions is computed as the distance between the centroids (average coordinate) of the set of vertices in each region. We used the midthickness surface, named fs LR.32k.L.midthickness.surf.gii, to attain the coordinates of each vertex. An exponential function was then fitted to the relationship between Euclidean distance and structural connectivity of each pair of regions in the parcellated connectome. The fast-conducting non-local projections (FNPs) were identified as pairs that were 3 standard deviations of log residuals above the exponential fit. **B.** FNPs identified in **A** are visualized as green lines on the cortical mesh. **C.** The evoked response of the cortical mesh to an impulse stimulus at the primary visual area, V1, is simulated and shown over time for three modulation strengths: **i.** *g* = 0.005, **ii.** *g* = 0.01, and **iii.** *g* = 0.02 s.

While sufficient modulation strengths are needed to amplify the functional relevance of FNPs, Fig. 4C shows that *C*(*t*) decays to zero more rapidly at strength *g*= 0.03 s than at *g* = 0.02 s, indicating that the FNP’s perturbation of dynamics is more persistent over longer timescales (*t >* 100 ms) at *g* = 0.02 s than at *g* = 0.03 s, i.e., intermediate gain rather than extreme values. This observation can be explained by examining the spatial patterns of the model’s evoked response under geometric and perturbed connectivity. For *g* = 0.02 s, the traveling wave response under geometric connectivity settles to a spatially nonuniform pattern that decays with distance from the stimulus (Fig. 4A(ii)), while the response under perturbed connectivity reaches a spatially uniform activation across all nodes (Fig. 4B(ii)). By contrast, for *g* = 0.03 s, the high modulation strength drives the response under geometric connectivity to fill the entire spatial extent (Fig. 4A(iii)), rendering its spatial pattern over longer timescales identical to that of the response under perturbed connectivity (Fig. 4B(iii)). This comparison suggests a non-monotonic relationship between modulation strength and the perturbation’s persistence over long timescales.

To quantify this relationship across the full spectrum of modulation strengths shown in Fig. 5A, we quantified the cosine dissimilarity between the zero-frequency components of the geometric and perturbed responses, denoted by *C_z_* (Eq. (**??**) in S3 Appendix) [64]. The zero-frequency component of each response is a linear approximation of the response when resolved over slow, order-of-seconds timescales by modalities such as functional magnetic resonance imaging (fMRI) [65]. Figure 5B shows that *C_z_* remains close to zero for *g* ⪅ 0.015 s, increases sharply to a maximum value after crossing this threshold, before decaying back to zero as *g* increases further. This non-monotonic behavior of *C_z_* indicates that FNP perturbations are most persistent over long timescales within a narrow range of modulation strengths, beyond which the perturbation becomes increasingly restricted to shorter timescales *t <* 100 ms.

Finally, to test whether these findings depend on the idealised ring geometry, we repeated the FNP experiment using empirical human connectivity. We fit the EDR to the Schaefer-400 left-hemisphere parcellated connectome (Fig. 6A), and identified FNPs as those region pairs lying more than three standard deviations of log residual above the fit, which together form a set of long-range projections linking posterior and anterior cortical regions (green, Fig. 6B). We then simulated the evoked response to an impulse stimulus applied to the primary visual cortex under two connectivity configurations — geometric connectivity alone (Fig. 6C) and the same connectivity with the empirical FNPs added (Fig. 6D) — across three modulation strengths.

The results recapitulate those obtained on the ring (Fig. 4). At low modulation (g = 0.005 s), the evoked response is weak and remains confined to regions proximal to the stimulus under both configurations, with the FNPs producing no discernible effect on the spatiotemporal pattern (Fig. 6C). As modulation strength increases, the two configurations diverge: by *t* = 100 ms the FNP-perturbed dynamics show activation in anterior regions that are not yet engaged under geometric connectivity alone, and which correspond to the distal endpoints of FNPs rather than being reached by progressive spread from the stimulus (Fig. 6D). Peak activation is essentially unchanged by the addition of FNPs at every modulation strength (*ϕ*_max_ = 228.6 *s^−^*^1^ versus 223.7 *s^−^*^1^, 342.2 *s^−^*^1^ versus 341.8 *s^−^*^1^, and 388.7 *s^−^*^1^ versus 390.8 *s^−^*^1^), confirming that FNPs reshape the spatial distribution of the evoked response rather than amplifying it. Thus the threshold-like dependence of FNP-mediated propagation on modulation strength, and the capacity of modulation to recruit specific long-range projections, both hold under anatomically realistic geometry.

### Effect of heterogeneous neuromodulation in the presence of long-range connections

The previous results demonstrate two mechanisms that can shape spatial patterns of traveling wave-evoked responses under modulation: (i) assigning spatial specificity to the modulation introduces anisotropy to the traveling wave; and (ii) adding FNPs to the structural connectivity enables additional faster propagation between remote populations (beyond what would result from geometric connectivity alone). Here we investigate whether, by leveraging both mechanisms, activity could propagate over long distances by modulating only the endpoints of a single FNP rather than the entire network. Such a selective routing mechanism would allow activity to reach its intended remote target without spreading to intermediate regions, providing both metabolic efficiency and communication specificity that could support adaptive cognition.

To isolate these mechanisms explicitly and disassociate their contributions from more realistic complex brain connectivity geometries, we again return to a ring topology. First, we tested whether modulating only two nodes of the ring network was sufficient to drive activity propagation between them, and then tested the impact of connecting them with an FNP in addition to the modulation. Figure 7A illustrates the evoked response under three conditions: (i) no modulation; (ii) both nodes modulated with strength *g* = 0.05 s; and (iii) both nodes are similarly modulated and additionally connected by an FNP. Figure 7A(ii) shows that while modulation amplifies the activity at the stimulus node and proximate nodes (relative to Fig. 7A(i)), at this strength it is insufficient to evoke a sustained response at the target node. By contrast, Fig. 7A(iii) shows that combining the same modulation with an FNP successfully drives sustained activation at the target and its proximate nodes - eliciting a traveling wave of evoked responses at the FNP end point in addition to the stimulus target. Together, these experiments demonstrate the critical role that FNPs play in enabling long-range propagation in the presence of flexible neuromodulatory tone.

**Fig 7.**
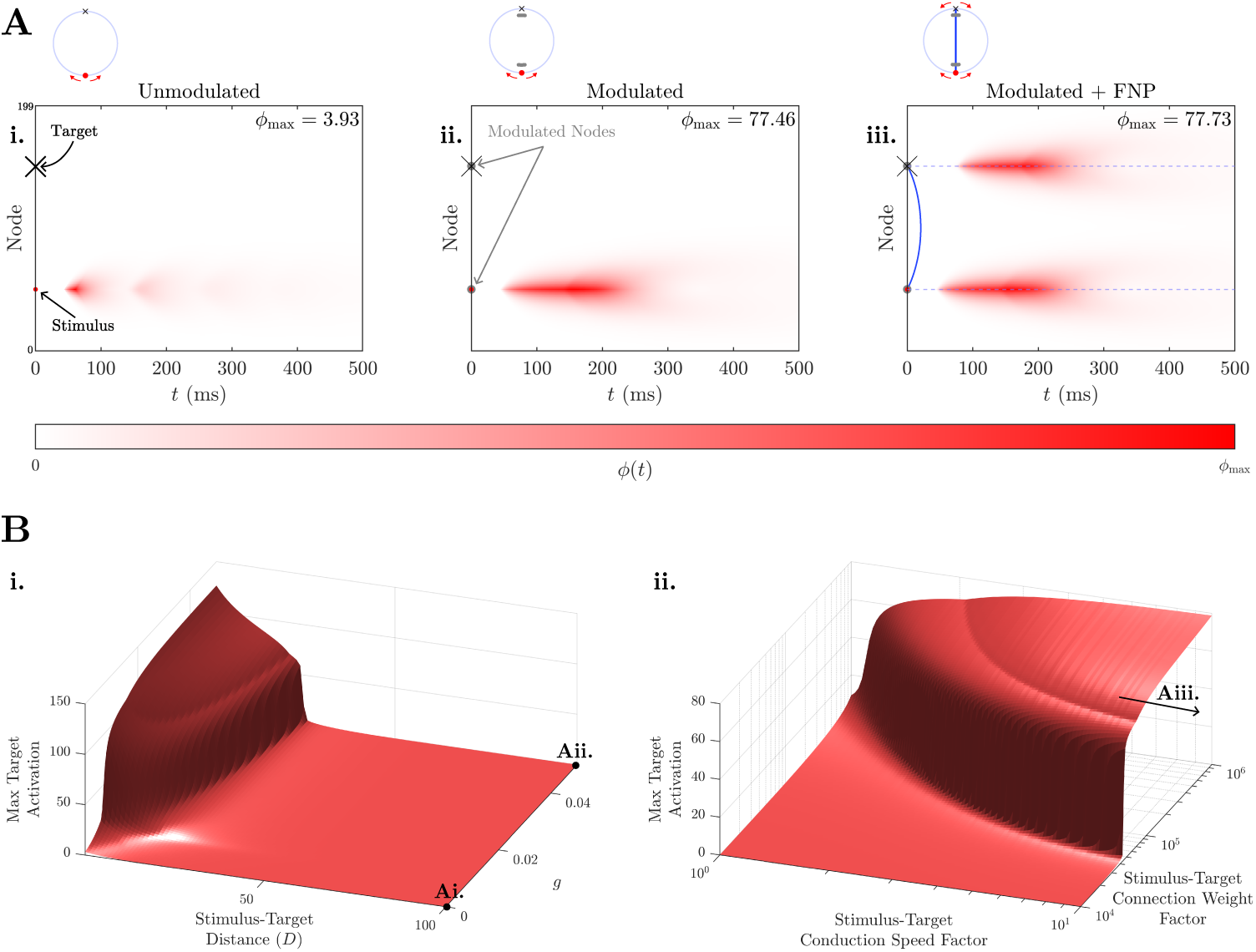
Combining spatially precise modulation with FNPs enables selective long-distance routing. **A.** The evoked response to a stimulus (node 50) was simulated under three conditions: **i.** No modulation or FNP; **ii.** Stimulus node and target node (node 150) modulated with strength *g* = 0.05 s; and **iii.** Stimulus and target nodes modulated at *g* = 0.05 s, while also being connected by an FNP (with same strength and conduction time used in previous experiments). **B. i.** The peak activation of the target node (maximum attained activity over time) in the absence of the FNP is plotted as a function of its distance from the stimulus node along the ring *D* and the strength of modulation applied to both nodes *g*. The parameter combinations considered in **A(i)**and **(ii)** are annotated. **ii.** The peak activation of the target node, when connected to the stimulus node by an FNP, is plotted as a function of the FNP’s enhanced strength (relative to geometric connectivity strength) and conduction velocity (relative to baseline conduction velocity), when the modulation strength applied to both nodes is fixed to *g* = 0.05 s. The parameter combination considered in **A(iii)** is denoted as an arrow (velocity *→ ∞*).

To better understand why an FNP was necessary to enable propagation between the two modulated nodes, we first investigated why geometric connectivity alone was insufficient to facilitate a sustained activation of the target node, as seen in Figs 7A(i) and (ii). We first swept the modulation strength and tracked the maximum firing rate of each node on the ring as a function of their distance to the stimulus site. Figure 7B(i) shows evoked responses decrease as a function of distance: *D*, (in units of nodes, with *D* = 1 corresponding to the stimulated node); and increase as a function of modulation strength *g*. This is expected since pairwise geometric connectivity reduces with distance, and modulation amplifies driving inputs. Interestingly, Fig. 7B(i) also shows a clear distance boundary in the *D − g* parameter space, where geometric connectivity can only facilitate stimulus wave propagation to a bounded distance over a broad range of neuromodulatory gain. By plotting this boundary across a much wider range of *g* values (see S3 Fig) we find that at the maximum distance *D* = 100, i.e., the desired stimulus target, the threshold neuromodulatory strength to propagate the stimulus wave was *g ≈* 0.75 s. This explains why *g* = 0 and *g* = 0.05 s in Fig. 7A(i) and (ii) resulted in no sustained activation at the target, as the neuromodulatory tone and the back-bone of geometric connectivity place the target node outside of this evoked response boundary.

Next, we examined what FNP properties enable the long-distance stimulus wave propagation seen in Fig. 7A(iii). Specifically, we characterize the contributions of FNP connection strength and conduction velocity when neuromodulatory tone is kept fixed (*g* = 0.05 s). Figure 7B(ii) shows that, similar to Fig. 7B(i), FNP parameters define a boundary separating a weak and strong target evoked response to the stimulus. It also allows us to contextualize our finding in Fig. 7A(iii)—indicated by the arrow on the surface—whereby those FNP properties place the FNP target in a strong evoked response regime and reduces the required modulation strength from *g ≈* 0.75 s (for geometric connectivity alone) to *g ≤* 0.05 s.

Together our findings suggests that the formation of FNPs, despite their developmental and metabolic costs, could play a key role in enabling otherwise metabolically expensive long-distance selective routing with physiologically plausible levels of modulation.

## Discussion

The cortex shows substantial spatiotemporal variability in its evoked responses to repeated presentations of the same stimulus, ranging from weak localized responses to strong spatially sustained responses of varying propagation directions. This work provides a biophysical account of a dynamic neuromodulatory mechanism that can give rise to this variability, while evolving on a static structural connectome. Our model demonstrates that even with a spatially homogeneous and isotropic (geometric) structural connectivity rule (which captures much of overall corticocortical connectivity), anisotropic activity propagation can be achieved with spatially heterogeneous modulation. Furthermore, our results reconcile the unique contributions of geometric and fast long-range connectivity to this propagation, showing how these contributions are modulated by time-varying dynamics from neuromodulatory systems. Taken together, this work provides a putative mechanism through which the brain may leverage unique aspects of its physical connectivity, allowing for the formation of privileged streams of activity propagation.

This work contributes to an emerging paradigm combining insights in corticocortical interactions with subcortical neurobiological mechanisms that can shape and constrain their on-going spatiotemporal properties [33, 42, 67–69]. Historically, a corticocentric approach has revealed how macroscale interactions across the cerebral cortex relate to structural networks via white-matter fiber connectivity [27, 70–72]. In parallel, an evolving body of work has focused on subcortico-cortical interactions, i.e., via the thalamus and brainstem, as a primary driver of on-going cortical dynamics [68, 73]. Our results combine these distinct perspectives, revealing how both elements of brain structure contribute to spatiotemporal activity propagation across the cerebral cortex. In particular, it shows how the significance of long-range corticocortical connectivity is regulated by neuromodulatory signaling from the brainstem, allowing specific aspects of static structure to be dominant over others and this dominance to vary in time.

We showed that long-range connections (or FNPs) have a minimum modulation threshold at which activity can propagate through. This threshold nonlinearity bears striking similarity with ignition, the empirical phenomenon that prefrontal cortical areas tend to activate in an all-or-none fashion in response to sensory stimuli [74]. While previous mechanistic explanations attribute ignition to nonlinear interactions between the stimulus and spontaneous cortical activity prior to stimulus onset [3, 29, 74, 75] - often incorporating a biologically unspecified vigilance signal [29] - our model positions the ascending neuromodulation as a neurobiological mechanism distinct from the cortex that may contribute to ignition phenomena. This suggests that both the neuromodulatory tone and spontaneous cortical activity may simultaneously gate ignition and poses several interesting questions about which mechanism predominates in different contexts, providing a key future direction for work linking theory to empirical observations of response variability.

The threshold nonlinearity of FNP propagation also demonstrates the importance of modulation in supporting the dynamical relevance of FNPs. Recent analyses have indicated that the dynamical properties of macroscale spontaneous (resting-state) cortical activity can be well captured by a simple distance-dependent or geometric connectivity rule that neglects the specificity of FNPs [19, 72, 76, 77]. The results of this work suggest that one possible reason contributing to the aforementioned successes of geometric models is that FNPs require sufficient modulation to magnify their dynamical effects to observable levels, and this modulation is seldom attained in spontaneous settings. However, we also found in Fig. 5 that the long-timescale perturbation induced by the FNP on cortical dynamics decreases with modulation strength beyond this threshold. This nonlinear relationship between long-timescale perturbation and modulation strength suggests that previous discrepant hypotheses on the timescales over which FNPs influence dynamics may be modulation-strength dependent; namely that for modulation strengths near the critical threshold, the role is maximized over all timescales [59, 78], and for modulation strengths exceeding the threshold, the role becomes increasingly restricted to shorter, millisecond timescales [64]. Thus, the level of neuromodulation may be a fundamental determinant of whether and how long FNPs shape cortical dynamics.

Beyond spatially diffuse modulation, our model predicts that sparse modulation of specific cortical regions can selectively promote propagation of activity between those regions, enabling targeted routing of activity over both short and long distances [42]. In this way, particular activity streams can be privileged over others in a context-dependent manner, which may be vital for adaptive cognition. While we manually specified which nodes to modulate in our simulations, anatomical evidence from the noradrenergic system suggests this capacity can be achieved in the physical brain by activating the numerous identified LC neurons whose axon collaterals have been found to project to sparsely distributed sets of cortical targets [62, 63].

Interestingly, a recent study reveals that the cortical targets of these sparsely projecting LC neurons exhibit functional correlations [79], consistent with the aforementioned hypothesis that these sparsely projecting LC neurons play an important role in privileging specific pairwise interactions.

The investigations in this work focused particularly on the modulatory effects of LC projections on excitatory populations in the cortex, as the internode projections that govern spatial propagation of activity in our model are excitatory. However, the cortical targets of LC neurons, given their diffuseness, are not confined to a specific neuronal type [53]. Furthermore, while we assumed that modulation results in a positive change in the gain of target populations (*g >*0), empirical experiments suggest that certain classes of adrenoceptors can have a negative effect [51, 80]. For example, it has been hypothesized that the *α*_2_ adrenoceptor can silence inhibitory neurons, leading to an excitation of total activity via disinhibition [81]. Future work could consider assigning differing modulation strengths (positive or negative) and adrenoceptor activation timescales to both excitatory and inhibitory populations, and investigate their consequent effects on evoked cortical dynamics. Another direction is to investigate the effects of modulating thalamic populations, given the large inputs they receive from neuromodulatory systems including the LC [53, 82]. Since the thalamic populations in our model project to cortical populations in the same node, it is likely that modulating them will alter the dominant temporal frequencies of the evoked response, but introducing more realistic thalamocortical heterogeneity into the model may also allow diverse spatial effects [83].

Our model experiments focused on exclusive phasic and tonic conditions of the neuromodulatory population so as to decompose their unique contributions to spatiotemporal stimulus evoked responses. In vivo recordings of the noradrenergic LC show these conditions are not mutually exclusive and instead LC activity shows a more continuous distribution of firing rates [56, 57]. However, by isolating tonic modulatory contributions we find these shift the steady-state operating point of the system towards an instability boundary, which predicts that marginal effects of phasic (burst) modulation on spatial stimulus propagation should diminish at elevated tonic baseline. This constitutes a proposed dynamical account of the inverted-U relationship posited by the adaptive gain theory [56]

In the present model we quantified neuromodulation as the change in excitatory population gain, defined as the slope of the population’s activation function about steady state. However, neuromodulation can be defined more generally between pairs of populations, as the change in the synaptic strength parameters *{ν_ab_}*. The benefit of this generalization is that it allows feedback pathways (*r → s, e → s, e → r*) and feedforward pathways (*s → e, s → i, s → r*) to be modulated independently [84]. With this ability, this model can be used to shed light on how the emphasis on internal prior predictions (via feedback modulation) versus external attention (via feedforward modulation) in the face of different stimuli is reflected quantitatively in the corresponding evoked-response dynamics [85]. This balance may be of importance in understanding the relationship between modulation dynamics and more subtle features of sensory stimuli that are not easily quantifiable; for example, the novelty of a stimulus in comparison to its precedents in a sequence [86].

The central motivation of this work was to put forward patterned neuromodulatory gain control as a possible neurobiological mechanism underpinning trial-to-trial variability in evoked cortical activity patterns, with the noradrenergic system as its best-characterized instantiation. The brain consists of many subcortical systems that project to the cortex in a relatively diffuse fashion [42, 83, 87], each involving a unique neurotransmitter, and the mechanism we describe is in principle available to each of them.

The model formalism developed here is not specific to any one neurotransmitter: gain modulation enters through a single timescale *η* and a set of population-specific strengths *g_a_*, and any diffusely projecting modulatory system could in principle be represented this way. What is noradrenergic in our implementation is the parameterisation rather than the formalism. Four choices carry this specificity. First, the modulation timescale *η* = 20 s*^−^*^1^ is set from adrenoceptor activation kinetics [52], and it is this timescale that determines the temporal resolution at which modulation can gate wave onset. Second, the stimulus-locked phasic firing mode is a characteristic LC input–output signature [35, 54]. Third, the coexistence of phasic and tonic regimes, and the behavioural significance attached to each, follows the adaptive gain account of LC function [56, 57]. Fourth, the heterogeneous single-neuron projection profiles that motivate our spatially sparse modulation are established for LC neurons specifically [62, 63].

Each of these would need revision for another ascending system, in ways that make the systems distinguishable in principle. Cholinergic modulation acts through both fast ionotropic and slow metabotropic receptors [80, 87], and so would require at least two modulation timescales rather than the single *η* used here, with correspondingly richer gating dynamics. Dopaminergic innervation of the cortex, by contrast, is markedly non-uniform: density follows cytoarchitectonic and hierarchical gradients rather than being distributed evenly [88–90]. This would convert the configurable modulated territories of Fig. 3B into anatomically constrained ones, and render the predicted anisotropy specific rather than arbitrary. Serotonergic receptor diversity includes both excitatory and inhibitory effects at the same target [91], which a single-signed *g* cannot represent and which would require the signed modulation discussed above; notably, 5-HT1A expression defines a distinct cortical gradient peaking in cingulate and default-mode regions [90].

This consideration applies in weaker form to the noradrenergic system itself. Noradrenaline transporter density is regionally graded across the human cortex [89], so the spatially uniform modulation strength we assume within modulated regions is an idealisation rather than a property of the system. Incorporating empirical receptor density maps as a spatially varying *g_j_* is a direct extension of the present model, and one for which parcellated human data are now openly available.

A further complication is that these systems do not act independently. While all arousal systems follow similar general principles of modulation, some project directly onto others; the locus coeruleus, for example, projects directly to the basal forebrain, the origin of the cholinergic system [92]. Arousal systems can also interact indirectly at overlapping cortical targets, where the joint concentrations of their released neurotransmitters exhibit nonlinear coupled dynamics [93, 94]. Each system’s influence on cortical dynamics therefore cannot be understood completely in isolation from the others. We accordingly present the contrasts above as structural differences that render the systems distinguishable in principle, rather than as demonstrated incompatibilities. Establishing which system is operative in a given context remains an empirical question, and one our model helps to pose.

Aside from arousal, the cortex also receives driving (and partially modulatory) inputs from diffusely projecting matrix populations in the thalamus [83], which differ from the core thalamic cells of our model which project strictly to the topographically aligned cortical region. Matrix projections can broadcast activity from the thalamus to multiple cortical regions simultaneously, and thus, have been hypothesized by both experimental and theoretical studies to support functional integration across large spatiotemporal scales necessary for conscious awareness [59, 67, 69, 95]. Future work could investigate the subtle differences in the dynamical effects of matrix thalamic inputs versus ascending modulator inputs, and the behavioral significance of these differences.

A key limitation of our model is that the input–output organization of neuromodulation was explored in the context of response variability, rather than determined from explicit biophysical constraints. Specifications included turning on the driving input from the external stimulus to generate phasic firing - simulating LC activation observed empirically - following relevant stimuli. While the mechanistic factors that control stimulus-dependent phasic activation remain incomplete [96], there is still scope for future modeling work to investigate how this activation may be influenced by a diverse range of inputs that ascending neuromodulatory systems receive, apart from sensory stimuli [53]. Of potential importance to the noradrenergic LC are the inputs from the prefrontal cortex, which have been hypothesized to control the switching of the LC between phasic and tonic firing behaviors [56, 97–99].

Additionally, whilst the present work focused on spatiotemporal neuromodulatory influences on stimulus evoked responses, there are likely many contributing mechanisms to this phenomenon - including global brain state changes not regulated by neuromodulation [100], shared input from other subcortical systems [101], and short-term plasticity effects most prevalent under repetitive stimulus protocols [102]. However, it still remains to be shown how these mechanisms integrate to form variable stimulus evoked responses and represents an exciting avenue for further research.

Other simplifying assumptions made here to aid mechanistic understanding include modeling the cortical sheet using a one-dimensional ring or three-dimensional surface geometry and assigning distance-dependent structural connectivity. Furthermore, aside from the last experiment, the conduction time of the FNP in our investigations was set to zero, under the assumption that it is negligible relative to the conduction times of projections under geometric connectivity. Despite being an idealization, this choice allowed us to model FNPs distinctly from EDR-predicted long-distance connections, without having to consider the uncertainty in empirical measurements of FNP conduction velocities [103]. Finally, the modulation strength in our experiments was assumed to be uniform across all nodes in the modulated region. More realistic spatial distributions of modulation targets can be modeled by the incorporation of receptor density maps [104], giving rise to further target-dependent effects of modulation.

Taken together, this work supports a view of the brain as a multiscale dynamical system in which arousal and thalamocortical architecture jointly regulate the selection and propagation of neural activity. Rather than being noise, variability in cortical responses reflects a structured, modulable feature of the system, and can rise from gain control and circuit heterogeneity. This perspective opens the door to a more general theory in which adaptive behavior emerges from the interplay between modulation, competition, and recurrent corticothalamic interactions.

## Author contributions

**Rishikesan Maran:** Conceptualization, Data curation, Formal analysis, Investigation, Methodology, Software, Validation, Visualization, Writing – original draft, Writing – review & editing.

**James M. Shine:** Writing – review & editing.

**Ben D. Fulcher:** Supervision, Project administration, Writing – review & editing. **Eli J. Müller:** Conceptualization, Data curation, Formal analysis, Funding acquisition, Investigation, Methodology, Project administration, Resources, Software, Supervision, Validation, Visualization, Writing – original draft, Writing – review & editing.

## Acknowledgments

R.M. would like to thank the Australian Government Research Training Program (RTP) Scholarship for financial support. B.D.F. acknowledges support from the Australian Research Council (FT240100418). E.J.M. acknowledges support from the Australian Research Council (DE250100540). The funders had no role in study design, data collection and analysis, decision to publish, or preparation of the manuscript.

## Supporting information

**S1 Fig.** Effect of tonic LC firing rate on stimulus-evoked responses. The model’s stimulus-evoked response over time is illustrated under tonic LC firing rates of **A.** *Q_l_* = 15 s*^−^*^1^, and **B.** *Q_l_* = 20 s*^−^*^1^. In both simulations, default geometric connectivity was used, node 100 was stimulated (shown by the red dot), and all nodes were modulated with strength *g* = 0.01 s.

**S2 Fig.** Effect of modulation delay on stimulus-evoked responses. The peak activation (maximum activity over time) of the stimulus-evoked response at each node 0 to 199, is illustrated over a finely resolved set of modulation delays (parameter *τ_lx_*). In all simulations, default geometric connectivity was used, node 100 was stimulated, and all nodes were modulated with strength *g* = 0.01 s.

**S3 Fig.** Threshold modulation for pairwise activation as a function of stimulus-target distance. The target activation, defined as the peak activation of the model’s stimulus-evoked response at a target node, was computed as a function of the *D*, the distance between the stimulus and target node along the ring, and *g*, the strength at which the stimulus and target node was modulated (all remaining nodes were not modulated). This function was first computed over parameter values *D* = 1, 2, *. . .* , 100 and *g* = 0, 0.001, 0.002, *. . .* , 1 s. A linear interpolation algorithm was then used to approximate contour plots of this function for function values 5, 10, 50 and 100 (i.e., when the peak activation at the target node is equal to 5, 10, 50 and 100 s*^−^*^1^).

**S1 Appendix. Corticothalamic model.** Equations of the corticothalamic neural field model and its extension to include a diffusely projecting neuromodulatory population.

**S2 Appendix. Numerical treatment of model.** Finite-difference schemes used to solve the model equations.

**S3 Appendix. Computation of cosine dissimilarity metrics.** Definitions of the cosine dissimilarity *C*(*t*) and its zero-frequency counterpart *C_z_*.

