## Supplementary Materials for "Dynamic Routing on a Static Structure: Neuromodulatory Gating of Cortical Wave Propagation"

**Contents**

S1 Fig. Effect of tonic LC firing rate on stimulus-evoked responses
S2 Fig. Effect of modulation delay on stimulus-evoked responses
S3 Fig. Threshold modulation for pairwise activation as a function of stimulus-target distance
S1 Appendix. Corticothalamic model
S2 Appendix. Numerical treatment of model
S3 Appendix. Computation of cosine dissimilarity metrics

### S1 Fig.

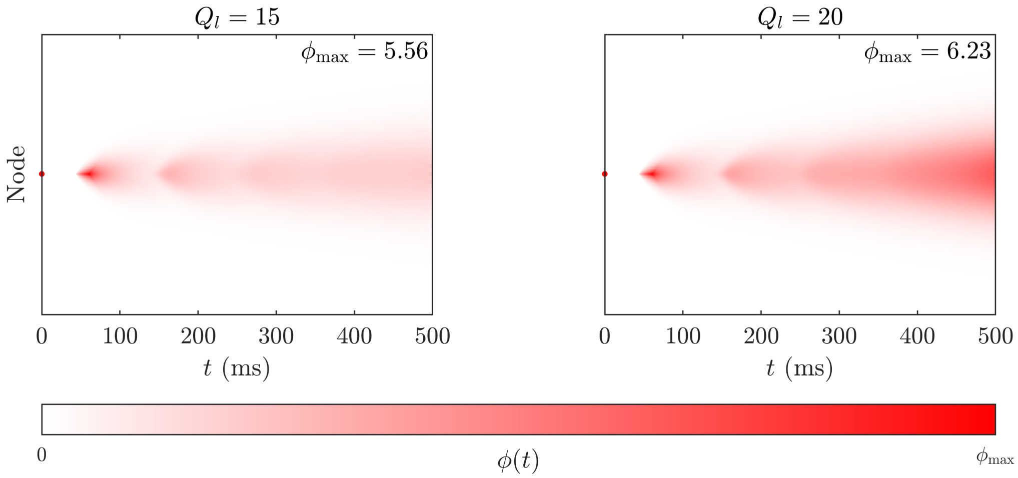


**Effect of tonic LC firing rate on stimulus-evoked responses** The model's stimulus-evoked response over time is illustrated under tonic LC firing rates of **A.** $Q_l = 15\ \mathrm{s}^{-1}$, and **B.** $Q_l = 20\ \mathrm{s}^{-1}$. In both simulations, default geometric connectivity was used, node 100 was stimulated (shown by the red dot), and all nodes were modulated with strength $g = 0.01\,\mathrm{s}$.

#### S2 Fig.

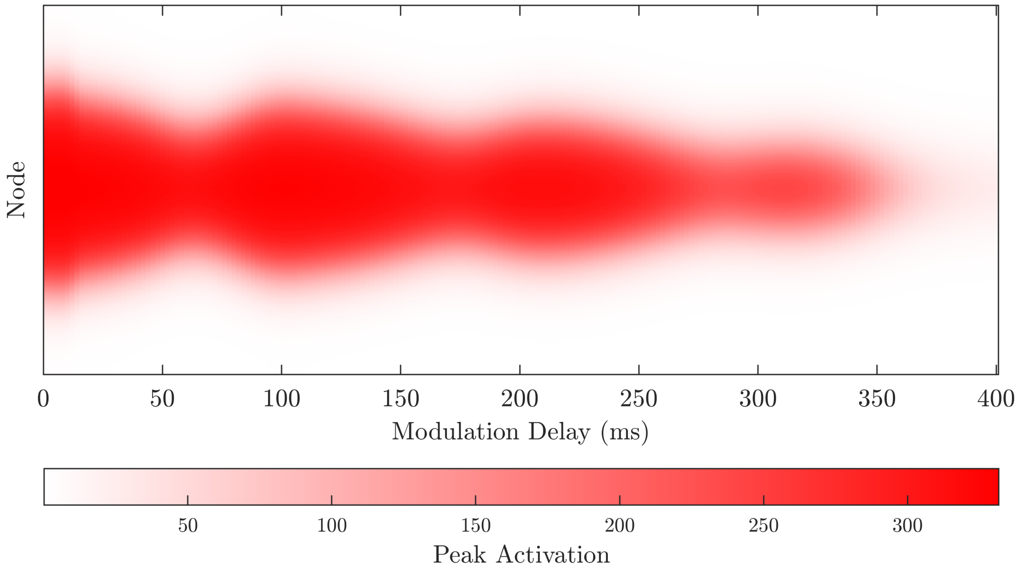


**Effect of modulation delay on stimulus-evoked responses** The peak activation (maximum activity over time) of the stimulus-evoked response at each node 0 to 199, is illustrated over a finely resolved set of modulation delays (parameter $\tau_{lx}$). In all simulations, default geometric connectivity was used, node 100 was stimulated, and all nodes were modulated with strength $g = 0.01\,\mathrm{s}$.

#### S3 Fig.

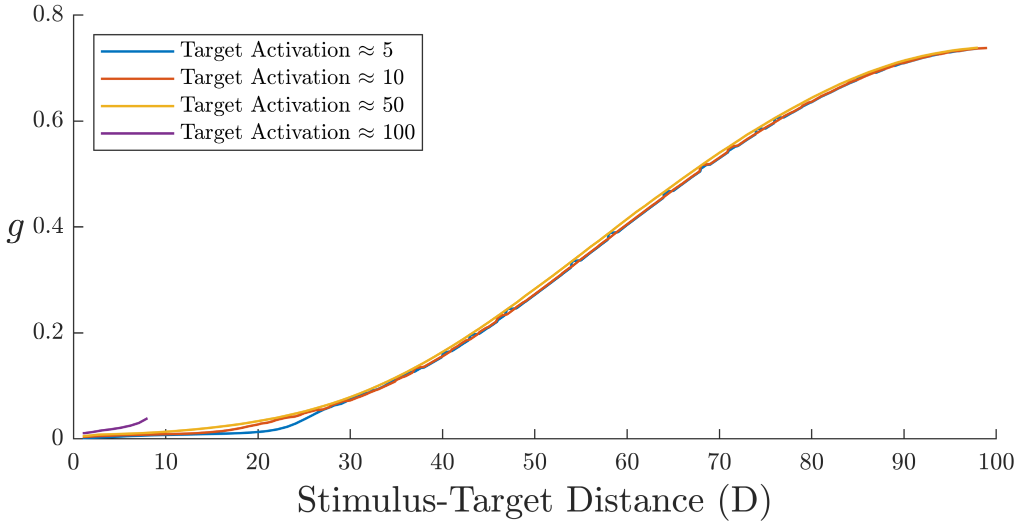


**Threshold modulation for pairwise activation as a function of stimulus-target distance.** The target activation, defined as the peak activation of the model's stimulus-evoked response at a target node, was computed as a function of the $D$, the distance between the stimulus and target node along the ring, and $g$, the strength at which the stimulus and target node was modulated (all remaining nodes were not modulated). This function was first computed over parameter values $D = 1, 2, \ldots, 100$ and $g = 0, 0.001, 0.002, \ldots, 1\,\mathrm{s}$. A linear interpolation algorithm was then used to approximate contour plots of this function for function values $5, 10, 50$ and $100$ (i.e., when the peak activation at the target node is equal to $5, 10, 50$ and $100\ \mathrm{s}^{-1}$).

### S1 Appendix. Corticothalamic model.

We describe the corticothalamic dynamic gain model used in this work, describing the relevant variables and mechanisms throughout. This model approximates the spatially distributed dynamics of macroscale (millimeter lengthscales and above) cortical and thalamic activity, under the influence of ascending neuromodulation.

**Existing Model Description.** The model is a network of $N$ coupled nodes, each representing a distinct cortical region and its topographically aligned thalamic region. Each node contains four distinct populations: excitatory ($e$) and inhibitory ($i$) neurons in the cortex, and reticular ($r$) and relay ($s$) nuclei in the thalamus. For each population $a = e, i, r, s$ at each node $j$, the model tracks over time $t$: the (mean) firing rate of generated pulses (spikes) outgoing from the somas of population $a$, $Q_a^j(t)$ in units $\mathrm{s}^{-1}$, and the (mean) membrane potential generated in the dendritic tree of population $a$, $V_a^j(t)$ in volts. Also, for each pair of populations $a, b$, the model tracks the (mean) rate of synaptic pulses at the the axon terminals of $b$, that target the dendrites of population $a$, $\phi_{ab}^j(t)$ in units $\mathrm{s}^{-1}$. The output variable of the model at each node is the self-excitatory propagator: $\phi^j(t) = \phi_{ee}^j(t),\ \forall j \in \{1, \ldots, N\}$.

The firing rate of each population is determined by its membrane potential in accordance to the sigmoidal activation function

$$Q_a^j(t) = S\left(V_a^j(t);\ Q_{\max}, \theta_a^j, \sigma_a^j\right) = \frac{Q_{\max}}{1 + \exp\left(-\frac{V_a^j(t) - \theta_a^j}{\sigma_a^j}\right)}\,. \tag{1}$$

This relationship is derived from the assumption that any given neuron in a population fires at its maximum firing rate $Q_{\max}$ if its membrane potential crosses a threshold [1], and the mean-field assumption that the threshold potentials across all neurons in the population follows a Gaussian-like distribution with mean $\theta_a^j$ and standard deviation $\sigma_a^j \pi/\sqrt{3}$ [2]. In the existing model, where modulation is not explicitly present, $\theta_a^j, \sigma_a^j$ are fixed parameters $\overline{\theta}, \overline{\sigma}$ over time, and equal across all populations and nodes.

At each node $j$, populations $i, r, s$ project to other populations in the same node in accordance to the equation

$$\phi_{ab}^j(t) = Q_b^j(t - \tau_{ab}),\ \forall a \in \{e, i, r, s\},\ b \in \{i, r, s\},\ j \in \{1, \ldots, N\}\,, \tag{2}$$

where $\tau_{ab}$ is the conduction time between the populations. If the projection is corticothalamic or thalamocortical, $\tau_{ab}$ is equal to the corticothalamic delay $t_0/2 > 0$, else $\tau_{ab} = 0$. Population $e$ additionally projects to populations in other nodes in accordance to the more general differential equation

$$\left(1 + \frac{1}{\gamma}\frac{d}{dt}\right)\phi_{ae}^j(t) = \sum_{k=1}^{N} \tilde{c}_{jk}\ Q_e^k(t - \tau_{ae} - d_{jk}/v_{jk}),\ \forall a \in \{e, i, r, s\}, j \in \{1, \ldots, N\}\,, \tag{3}$$

where $\gamma$ is the cortical damping rate, $\tilde{c}_{jk} = c_{jk}/\sum_j c_{jk}$ is the normalized connectivity strength of the corticocortical axonal projection from node $k$ to $j$, and $v_{jk}$ is its conduction speed. Equation (3) aims to capture the wave propagation dynamic of the original continuous-space neural field model, now on a discrete network topology. To justify Eq. (3), we equivalently express it as the (sum of) Volterra equations

$$\phi_{ae}^j(t) = \sum_{k=1}^{N} \int_{-\infty}^{t} dt'\, F_{jk}(t - t')\, Q_e^k(t' - \tau_{ae}), \text{ where} \tag{4}$$

$$F_{jk}(t) = \tilde{c}_{jk}\,\gamma\,\exp\left(-\gamma\,(t - d_{jk}/v_{jk})\right)\Theta(t - d_{jk}/v_{jk}) \tag{5}$$

is the response of node $j$ to an impulse at node $k$ ($Q_e^k(t' - \tau_{ae}) = \delta(t' - \tau_{ae})$), and $\Theta$ is the Heaviside function. In the default setting of the model, where the $N$ nodes are arranged in a ring, connectivity strengths are determined by an EDR ($\tilde{c}_{jk} \propto \exp(-\lambda d_{jk})$), and all conduction velocities are equal ($v_{jk} \equiv v$), $F_{jk}(t)$ captures four important properties of $G(t, \mathbf{r})$, the impulse response (Green's function) of the wave equation in the original neural field model [1]:

- The impulse response is zero up until time $d_{jk}/v$, reflecting the finite velocity of axonal propagation from node $k$ to $j$.
- The time-integrated impulse response of node $j$ is $\int_0^\infty F_{jk}(t)dt = \tilde{c}_{jk} \propto \exp(-\lambda d_{jk})$, which abides by the EDR;
- The aggregate impulse response across all nodes at time $t$ is

$$\sum_j F_{jk}(t) = \gamma \exp(-\gamma t) \frac{\sum_j \exp(-\lambda d_{jk}) \exp(\gamma d_{jk}/v) \Theta(t - d_{jk}/v)}{\sum_j \exp(-\lambda d_{jk})}, \tag{6}$$

  which is a function that is zero for time $t < 0$, then increases to a maximum before decaying back to zero at rate $\gamma$ as $t \to \infty$. This property aligns with the space-integrated aggregate impulse response in the original neural field model [1],

$$\int_{\mathbb{R}^2} d^2\mathbf{r}\ G(t, \mathbf{r}) = \gamma^2 t \exp(-\gamma t) \Theta(t), \tag{7}$$

  and this property ensures that the model is linearly stable at its physiologically plausible fixed points (c.f. the Fourier transform of this response, $(1 - i\omega/\gamma)^2$, in Equation 11 of [3]);
- The time-integrated aggregate impulse response across all nodes is $\sum_j \tilde{c}_{jk} = 1$, which is required to ensure conservation of pulses.

By abiding these properties, we argue that Eq. 3 sufficiently recapitulates the traveling waves produced by the wave equation in the original neural field model.

Following propagation between populations, incoming pulses at a population of a given node are converted into postsynaptic membrane potentials in accordance with

$$\left(1 + \frac{1}{\alpha} \frac{d}{dt}\right) \left(1 + \frac{1}{\beta} \frac{d}{dt}\right) V_a^j(t) = \sum_{b \in \{e,i,r,s\}} \nu_{ab}\ \phi_{ab}^j(t), \qquad \forall a \in \{e, i, r, s\},\ j \in \{1, \ldots, N\}, \tag{8}$$

where $\alpha, \beta$ are the rates at which the potential rises and falls, and $\nu_{ab}$ is the synaptic strength. If population $b$ is not synaptically coupled to $a$ per the node architecture in Fig 1B of the main text, then $\nu_{ab} = 0$. The membrane potential of the relay population $s$ in each node is also driven by the stimulus propagator $\phi_{sx}$ with synaptic strength $\nu_{sx}$.

The steady-state firing rates of each population, $\overline{Q_e}, \overline{Q_i}(= \overline{Q_e}), \overline{Q_r}, \overline{Q_s}$, are solved by setting all derivatives as zero, yielding the system of equations [3]:

$$\begin{cases} S^{-1}\left(\overline{Q_e}\right) &= (\nu_{ee} + \nu_{ei})\overline{Q_e} + \nu_{es}\overline{Q_s}, \\ S^{-1}\left(\overline{Q_r}\right) &= \nu_{rs}\overline{Q_s} + \nu_{re}\overline{Q_e}, \\ S^{-1}\left(\overline{Q_s}\right) &= \nu_{se}\overline{Q_e} + \nu_{sr}\overline{Q_r} + \nu_{sx}\overline{\phi_x}\ , \end{cases} \tag{9}$$

where $S^{-1}(\ \cdot\ )$ is the inverse function of $S(\ \cdot\ ; Q_{\max}, \overline{\theta}, \overline{\sigma})$, and $\overline{\phi_x}$ is the steady state component of $\phi_{sx}(t)$ across all nodes. After numerically solving this system, the steady-state membrane potentials and synaptic pulse rates are then computed as $\overline{V_a} = S^{-1}\left(\overline{Q_a}\right)$, $\overline{\phi_{ab}} = \overline{Q_b}$. We then tracked variables $Q_a^j, V_a^j, \phi_{ab}^j$ (for all $a, b, j$) as perturbations from their steady states. From the equations above, only the nonlinear equation Eq. (1) is formulated differently under this redefinition:

$$Q_a^j(t) = S\left(\overline{V_a} + V_a^j(t);\ Q_{\max}, \theta_a^j, \sigma_a^j\right) - \overline{Q_a},\ \forall a \in \{e, i, r, s\},\ j \in \{1, \ldots, N\}\,. \quad (10)$$

The model's stimulus-evoked response is treated as perturbations of the output, $\{\phi_{ee}^j(t)\}_j$, about its steady state, when the stimulus propagator at a single node $j_0$ is perturbed with a Delta function:

$$\phi_{sx}^{j_0}(t) = \delta(t - t_{os})\,, \quad (11)$$

where $t_{os} > 0$ is the stimulus onset time.

**Modeling Neuromodulation of Stimulus-Evoked Responses.** We introduce a neuromodulatory population (NM), indexed $l$, that is separate from the existing $N$ nodes. The model tracks the mean membrane potential of the NM $V_l(t)$, its mean firing rate $Q_l(t)$, and the mean synaptic pulse rate targeting the populations in each node, $\phi_{al}^j(t)$ for $a \in \{e, i, r, s\},\ j \in \{1, .., N\}$, all as perturbations from their respective steady states.

The firing rate of the NM population is a sum of a phasic and tonic component. The tonic component is a constant value ($Q_l^{\text{to}}(t) \equiv Q_l^{\text{to}}$). The phasic component is determined by the membrane potentials driven by the stimulus propagator $\phi_{sx}$:

$$\left(1 + \frac{1}{\alpha}\frac{d}{dt}\right)\left(1 + \frac{1}{\beta}\frac{d}{dt}\right) V_l^{\text{ph}}(t) = \nu_{lx}\ \phi_{sx}^{j_0}(t - \tau_{lx})\,, \quad (12)$$

where $j_0$ is the stimulated node and $\tau_{lx}$ is a time delay. By default, we assumed that NM activation is time-locked to the stimulus at the target node ($\tau_{lx} = 0$), however as part of our investigations we also consider a lagged relationship ($\tau_{lx} > 0$). The membrane potential is then related to the phasic firing rate by

$$Q_l^{\text{ph}}(t) = \rho_l V_l^{\text{ph}}(t)\,, \quad (13)$$

where $\rho_l$ is the gain of the NM. An important distinction to note is that Eq. (13) is a linearization of the NM population's nonlinear sigmoidal activation function about its steady state. This is because computing the NM firing rate from $V_l^{\text{ph}}$ via the sigmoidal relationship in Eq. (10) requires its steady-state membrane potential ($\overline{V_l^{\text{ph}}}$) which, unlike the other populations, cannot be computed under our current model formulation.

The incoming pulse rate to the cortex or thalamus is computed from the firing rate in a similar manner to populations $i, r, s$:

$$\phi_{al}^j(t) = Q_l^{\text{to}} + Q_l^{\text{ph}}(t - \tau_{al}),\ \forall a \in \{e, i, r, s\},\ j \in \{1, \ldots, N\}\,, \quad (14)$$

where $\tau_{al}$ is the conduction time from the NM to population $a$.

For each population $a$ in node $j$, we define $\overline{\rho_a}$ as the steady-state gain of population $a$, and $\rho_a^j(t)$ the dynamic perturbation of this gain from steady state under the influence of modulation. We also define the modulation factor as the dynamic gain divided by its steady state, and $M_a^j(t)$, the perturbation in modulation factor from its steady-state value of 1. Incoming pulses then drive the modulation factor with timescale $\eta$ and strength $g_a^j$:

$$\left(1 + \frac{1}{\eta}\frac{d}{dt}\right) M_a^j(t) = g_a^j \phi_{al}(t),\ \forall a \in \{e, i, r, s\},\ j \in \{1, .., N\}\,. \quad (15)$$

Changes in $M_a^j(t)$ then drive proportional changes in population gain, in accordance to the definition $\rho_a^j(t) = \overline{\rho_a} M_a^j(t)$. Since the gain is not an explicit parameter of the sigmoidal activation function, it is adjusted indirectly by adjusting the mean firing threshold and threshold spread of each population in each node $(\theta_a^j, \sigma_a^j)$. To do this, we note that the gain of population $a$ is

$$\overline{\rho_a} + \rho_a^j(t) = \left[\frac{dQ_a^j}{dV_a^j}\right]_{V=\overline{V_a}} = \frac{\overline{Q_a}}{\sigma_a^j}\left(1 - \frac{\overline{Q_a}}{Q_{\max}}\right).$$

Under a fixed $Q_{\max}, \overline{Q_a}$, the gain is inversely proportional to $\sigma_a^j$. Hence to change the gain by $\overline{\rho_a} M_a^j(t)$, the total gain is multiplied by $1 + M_a^j(t)$, which is achieved by setting $\sigma_a^j, \theta_a^j$ as.

$$\sigma_a^j(t) = \frac{\overline{\sigma}}{1 + M_a^j(t)}; \; \frac{\overline{V_a} - \theta_a^j(t)}{\sigma_a^j(t)} = \frac{\overline{V_a} - \overline{\theta}}{\overline{\sigma}}, \; \forall a \in \{e, i, r, s\}, \; j \in \{1, .., N\}. \quad (16)$$

The second equation determines the value of $\theta_a^j$ that keeps the steady state $(\overline{V_a}, \overline{Q_a})$ unchanged.

### S2 Appendix. Numerical treatment of model.

The equations of the model described in S1 Appendix were numerically solved using a finite difference scheme. The time domain was truncated to a bounded interval $[0, T]$, from which we partitioned time points $\{t_n = n\Delta t : n = 0, \dots, N_t\}$ with time step $\Delta t = T/N_t$. We define $Q_a^j[n], V_a^j[n], \phi_{ab}^j[n], Q_l^{\text{ph}}[n], M_a^j[n], \theta_a^j[n], \sigma_a^j[n]$ as the numerical solutions at each node $j$ and time step $t_n$.

We first computed the steady states $\overline{V_a}, \overline{Q_a}$ of the model. This was done by numerically solving the system of equations in Eq. (9) via the `fzero` function in MATLAB.

We assumed the trivial initial condition that activity is initially at steady state and unmodulated — hence $Q_a^j[n] = V_a^j[n] = \phi_{ab}^j[n] = M_a^j[n] = 0$ for all $a, b, j$ and time steps $n \leq 0$. Then for $n \geq 1$, we numerically solved Eqs (2), (3), (8), (10), (12), (13), (14), (15), (16) using finite differences. We describe these computations below in the same order as the scheme:

$V_a^j[n]$ is computed from $V_a^j[n-1], V_a^j[n-2]$ and $\phi_{ab}^j[n-1]$ using the central difference approximation of Eq. (8):

$$\begin{aligned} V_a^j[n-1] + \left(\frac{1}{\alpha} + \frac{1}{\beta}\right) \frac{V_a^j[n] - V_a^j[n-2]}{2\Delta t} + \frac{1}{\alpha\beta} \frac{V_a^j[n] - 2V_a^j[n-1] + V_a^j[n-2]}{\Delta t^2}, \\ = \sum_{b \in \{e,i,r,s,x\}} \nu_{ab} \phi_{ab}^j[n-1], \ \forall a \in \{e, i, r, s\}. \end{aligned} \tag{17}$$

$Q_l^{\text{ph}}[n]$ is computed from $Q_l^{\text{ph}}[n-1], Q_l^{\text{ph}}[n-2]$ and $\phi_{sx}^{j_0}[n-1]$ ($j_0$ = stimulated node) by combining Eqs (12) and (13) and applying a central difference approximation:

$$\begin{aligned} Q_l^{\text{ph}}[n-1] + \left(\frac{1}{\alpha} + \frac{1}{\beta}\right) \frac{Q_l^{\text{ph}}[n] - Q_l^{\text{ph}}[n-2]}{2\Delta t} + \frac{1}{\alpha\beta} \frac{Q_l^{\text{ph}}[n] - 2Q_l^{\text{ph}}[n-1] + Q_l^{\text{ph}}[n-2]}{\Delta t^2}, \\ = \rho_l \nu_{lx} \phi_{sx}^{j_0}[n - 1 - [\tau_{lx}/\Delta t]], \end{aligned} \tag{18}$$

where $[\tau_{lx}/\Delta t]$ is $\tau_{lx}/\Delta t$ rounded to the nearest integer.

$M_a^j[n]$ is computed from $M_a^j[n-1], Q_l^{\text{to}}$ and $Q_l^{\text{ph}}[n]$ by combining Eqs (14) and (15) and using a Backward Euler Scheme [4]:

$$M_a^j[n] + \frac{M_a^j[n] - M_a^j[n-1]}{\eta\Delta t} = g_a^j(Q_l^{\text{to}} + Q_l^{\text{ph}}[n - [\tau_{al}/\Delta t]]). \tag{19}$$

The firing threshold mean and spread $\theta_a^j[n], \sigma_a^j[n]$ are computed from $M_a^j[n]$ using Eq. (16):

$$\sigma_a^j[n] = \frac{\overline{\sigma}}{1 + M_a^j[n]}, \ \frac{\overline{V_a} - \theta_a^j[n]}{\sigma_a^j[n]} = \frac{\overline{V_a} - \overline{\theta}}{\overline{\sigma}}. \tag{20}$$

$Q_a^j[n]$ is computed from $V_a^j, \theta_a^j[n]$ and $\sigma_a^j[n]$ using Eq. (10):

$$Q_a^j[n] = S\left(\overline{V_a} + V_a^j[n];\ Q_{\max}, \theta_a^j[n], \sigma_a^j[n]\right) - \overline{Q_a}, \ \forall a \in \{e, i, r, s\}. \tag{21}$$

Since $\overline{V_a}, \overline{Q_a}$ are solved numerically from Eq. (9), their numerical errors will carry on to $Q_a^j[n]$ via Eq. (21). This means when $V_a^j[n]$ is zero, computing $Q_a^j[n]$ from Eq. (21) will return a non-zero value $\epsilon$. This error will also be amplified by the modulation factor $M_a^j$, since $Q_a^j(t) = \overline{\rho_a}(1 + M_a^j)V_a^j(t) + o(V_a^j(t)^2)$. To correct this error, we set $Q_a^j[n] = 0$ if $\overline{\rho_a}|V_a^j[n]| \leq \epsilon$.

For populations $b = i, r, s$, $\phi^j_{ab}[n]$ is computed from $Q^j_b[n]$ using Eq. (2):

$$\phi^j_{ab}[n] = Q^j_b[n - [\tau_{ab}/\Delta t]] \,. \tag{22}$$

For the excitatory population $e$, $\phi^j_{ae}[n]$ is computed using Eq. (3). For accessibility, we first rewrite this equation as

$$\left(1 + \frac{1}{\gamma}\frac{d}{dt}\right)\phi^j_{ae}(t) = P^j_e(t - \tau_{ae}), \text{ where } P^j_e(t) = \sum_{k=1}^{N} \tilde{c}_{jk}\ Q^j_e(t - d_{jk}/v_{jk}) \,. \tag{23}$$

We then computed $P^j_e[n]$ and applied a Backward Euler Scheme:

$$P^j_e[n] = \sum_{k=1}^{n} \tilde{c}_{jk}\ Q^j_e[n - [(d_{jk}/v_{jk})\,/\Delta t]] \,; \tag{24}$$

$$\phi^j_{ae}[n] + \frac{\phi^j_{ae}[n] - \phi^j_{ae}[n-1]}{\gamma \Delta t} = P^j_e[n - [\tau_{ae}/\Delta t]]. \tag{25}$$

Finally, we computed the stimulus propagator $\phi^{j_0}_{sx}(t) = \delta(t - t_{os})$. This delta functional was numerically computed with a Gaussian mollifer for smoothness:

$$\phi^{j_0}_{sx}[n] \propto \exp\left(-\frac{(t_n - t_{os})^2}{\sigma_t^2}\right), \text{ such that } \sum_{n=1}^{N_t} \phi^{j_0}_{sx}[n]\Delta t = 1 \,. \tag{26}$$

For the investigations reported here, we set $\Delta t = 2^{-14}$ s, a value which ensures numerical stability, and set $\sigma_t = \Delta t$.

#### S3 Appendix. Computation of cosine dissimilarity metrics.

The cosine dissimilarity between the model's stimulus-evoked dynamics under geometric connectivity and the dynamics under hybrid connectivity (containing additional FNPs) is defined by

$$C(t) = 1 - \frac{\sum_{j=1}^{N} \left( \phi_0^j(t)\ \phi_1^j(t) \right)}{\sqrt{\sum_{j=1}^{N} \left( \phi_0^j(t) \right)^2} \sqrt{\sum_{j=1}^{N} \left( \phi_1^j(t) \right)^2}}\,, \tag{27}$$

where $\phi_0^j$ and $\phi_1^j$ are model's output at node $j$ under geometric and hybrid connectivity respectively.

Similarly, the cosine dissimilarity between the zero-frequency components of the model's stimulus-evoked response under geometric and hybrid connectivity is defined by

$$C_z = 1 - \frac{\sum_{j=1}^{N} \left( z_0^j\ z_1^j \right)}{\sqrt{\sum_{j=1}^{N} \left( z_0^j \right)^2} \sqrt{\sum_{j=1}^{N} \left( z_1^j \right)^2}}\,, \tag{28}$$

where $z_0^j$ and $z_1^j$ are zero-frequency components of $\phi_0^j(t)$ and $\phi_1^j(t)$, respectively.
